# Adversity and adolescent brain development: differential associations with grey and white matter across two longitudinal cohorts

**DOI:** 10.64898/2026.07.06.736703

**Authors:** Lea C. Michel, Divyangana Rakesh, Tobias Banaschewski, Gareth J. Barker, Arun L.W. Bokde, Rüdiger Brühl, Sylvane Desrivières, Herta Flor, Penny Gowland, Antoine Grigis, Andreas Heinz, Herve Lemaitre, Frauke Nees, Dimitri Papadopoulos Orfanos, Tomáš Paus, Luise Poustka, Michael N. Smolka, Nathalie Holz, Nilakshi Vaidya, Henrik Walter, Robert Whelan, Paul Wirsching, Gunter Schumann, IMAGEN Consortium, Delia Fuhrmann, Rogier A. Kievit

## Abstract

Globally, 60% of the population has experienced at least one type of adversity (e.g., emotional abuse, bullying) across infancy, childhood, and adolescence. Such experiences have been linked to an increased risk for mental health disorders. Changes in brain structure following experiences of childhood adversity have been hypothesised to be a mechanistic pathway explaining later mental health issues. However, to understand how changes in brain structure might mediate the effects of adversity, it is essential to identify which underlying neuronal processes may be affected by different types of adverse experiences. A key open question is whether grey or white matter is more vulnerable to adversity, as these two structures reflect distinct neurobiological mechanisms.

This study investigated whether differences in trajectories of grey and white matter development during adolescence can be explained by exposure to different types of adversity. We applied the Adverse Adolescent Experiences Framework (Pollmann et al., 2025) categorising adversity into four levels: Intrapersonal (e.g., accidents), Caregiver (e.g., emotional neglect), Peer (e.g., bullying), and Community (e.g., neighbourhood safety). Exposure to each of the four factors was estimated through principal components analyses. We analysed two large longitudinal datasets: the Adolescent Brain Cognitive Development study (∼12,000 adolescents measured at ages 10, 12, and 14) and the IMAGEN study (∼1,400 adolescents measured at ages 14, 19, and 22). Using latent growth curve models, we captured individual differences in brain development by estimating baseline levels (intercepts) and rates of change (slopes) for total grey matter volume and mean white matter fractional anisotropy.

In both cohorts, we found significant interindividual variability in baseline levels and rates of change for both grey matter volume and fractional anisotropy. Caregiver, Peer, and Community adversities were negatively associated only with the intercepts of grey matter volume and white matter fractional anisotropy. Importantly, associations differed between grey and white matter. In ABCD, Peer and Community adversities were more strongly associated with grey matter volume intercepts. In contrast, in IMAGEN, Caregiver, Peer and Community adversities were more strongly linked to white matter fractional anisotropy intercepts. This suggests that adversity has unique associations with grey and white matter, rather than exerting a uniform influence on brain structure.

By demonstrating that different environments generate distinct biological associations with brain maturation, this work underscores the need to consider both grey and white matter when assessing the neurodevelopmental pathways to outcomes across the lifespan.

## Introduction

Adolescence is a time of heightened sensitivity to environmental influences, as adolescents navigate profound physical, social, and emotional transitions (Fuhrmann et al., 2015; Paus et al., 2008). These changes coincide with increased vulnerability to mental health difficulties (Blakemore, 2019; Sisk & Gee, 2022), with 48.4% of individuals with a mental disorder showing onset before 18 years of age (Solmi et al., 2022).

Mental health disorders are particularly more prevalent in adolescents who have experienced adversity (Kessler et al., 2010). The term adversity refers to potentially traumatic events defined as exposure to experiences that demand significant psychological, social, or neurobiological adaptation (Frankenhuis & Amir, 2022; McLaughlin, 2016), such as household dysfunction, emotional abuse, and bullying. Meta-analyses show that 60% of the population has experienced at least one adverse childhood event (Madigan et al., 2023), and that individuals who experienced multiple types of childhood adversity were at increased risk of later mental health issues (Hughes et al., 2017).

Adversity is also a key environmental factor associated with brain development (McLaughlin et al., 2019; Nelson & Gabard-Durnam, 2020). Both reports of individual adversities (Holz et al., 2023; Hong et al., 2021; Pollok et al., 2022) and cumulative exposure (Dannlowski et al., 2012; Paquola et al., 2016) are linked to differences in brain structure. Childhood and adolescence are identified as sensitive periods for brain maturation, characterised by changes across the whole cortex (Norbom et al., 2021; Lebel et al., 2019). Thus, understanding how adversity contribute to interindividual variability in brain development is crucial for explaining why some adolescents are more vulnerable to mental health problems (Holz et al., 2023; Pollok et al., 2022; Whittle et al., 2025).

One likely source of variation in brain development, as well as vulnerability to adversity, is biological sex. Sex differences have been observed in both grey and white matter development (Ingalhalikar et al., 2014; Kaczkurkin et al., 2019; although the literature remains divided, see Eliot et al., 2021) and in how individuals experience and are biologically affected by adversity (Bath, 2020; Peng et al., 2025; Shaul et al., 2026). Yet, few studies have investigated how male and female brain development are differently associated with adversity. Results in the literature suggest sex-specific associations (Samplin et al., 2013; Teicher et al., 2018) which underscore the need to examine sex differences as a fundamental source of neurodevelopmental variability to uncover mechanistic pathways following adversity.

To better comprehend how changes in brain structure serve as a mechanism for later mental health issues, we need a clearer understanding of the neuronal mechanisms associated with different adversities. To this end, it is essential to compare how different types of adversity relate to distinct features of neurodevelopment, such as grey and white matter structures. At the cellular level, grey and white matter are both composed of neurons, but include distinct components: grey matter primarily includes cell bodies, dendrites, and synapses, while white matter primarily consists of myelinated axons. Animal research indicates that chronic stress can result in cellular-level changes, including dendritic atrophy, synaptic loss, or altered myelination (Lehmann et al., 2017; Vyas et al., 2002; Watanabe et al., 1992). Hence, if adversity is differentially associated with distinct features of grey and white matter, this could provide critical insights into the specific neurobiological processes (e.g., myelination, synaptic pruning) that are affected by adverse experiences.

To date, few studies have examined how adversity is differentially linked with the development of cortical grey and white matter structures (Hong et al., 2021; Levesque et al., 2015; Sheridan et al., 2022; Thijssen et al., 2024; Zuo et al., 2019). These studies generally found that adversity predicted both grey and white matter structural metrics (Hong et al., 2021; Sheridan et al., 2022; Thijssen et al., 2024) or only one of those metrics (Levesque et al., 2015; Zuo et al., 2019). For instance, maternal hostile parenting (Levesque et al., 2015) was associated with differences in grey matter volume but not white matter volume in adolescence, which could reflect associations with distinct neuronal mechanisms. However, no study directly compared whether grey or white matter was more vulnerable to adversity, or whether they were differentially vulnerable to distinct components of adversity. In other fields, existing studies that examine both tissue types suggest they provide partially overlapping but also unique information in relation to behaviour (Michel et al., 2024), but also that their changes coincide during periods of accelerated change, such as ageing or neurodegenerative diseases (Agosta et al., 2011; Hoagey et al., 2025).

The present study addresses this gap by investigating whether exposure to different forms of adversity are associated with distinct patterns of grey and white matter development during adolescence. Adversity was conceptualised using the Adverse Adolescent Experiences Framework (Pollmann et al., 2025), which was developed to identify the unique challenges and experiences that characterise adolescence. The framework categorises adversity into four levels: Intrapersonal, Caregiver, Peer, and Community. Exposure to adversity in each domain was estimated using probabilistic principal component analyses (pPCA). For our grey and white matter metrics, we chose grey matter volume and fractional anisotropy, given their widespread use in the literature. Developmental trajectories of total grey matter volume and mean white matter fractional anisotropy were modelled using latent growth curve models, enabling estimation of both baseline levels (intercepts) and rates of change (slopes) for each individual. We examined how different measures of adversity predicted individual differences in grey and white matter levels and changes. To enhance generalisability, we leveraged two large, complementary, and independent longitudinal datasets: the Adolescent Brain Cognitive Development (ABCD) study (N = 11,876 at ages 10, 12, and 14; Volkow et al., 2018) and the IMAGEN study (N = 2,090 at ages 14, 19, and 22; Mascarell Maričić et al., 2020). This study aims to investigate how different types of adversity predict grey and white matter development, and test which metric, grey matter volume or fractional anisotropy, is more sensitive to adversity in our two datasets.

## Methods

### Participants

This study used data from ABCD and IMAGEN cohorts. The ABCD Study (https://abcdstudy.org/) is an ongoing longitudinal study conducted across 21 data acquisition sites in the United States, enrolling 11,876 adolescents aged 9 to 16 years. For more detailed information on ABCD protocols and inclusion/exclusion criteria, see Volkow et al. (2018). This paper analysed the three imaging time points from release 5.1 (https://abcdstudy.org/; https://nda.nih.gov/study.html?id=2313), from which we included 9,994 adolescents (4,904 males and 4,488 females) across the three time points (i.e., at ages 10, 12, and 14). We excluded MRI time points if the quality of the T1 or DWI scans was poor (688 T1 scans and 1,351 DWI scans) or if the absolute z-score of the neuroimaging metric of interest exceeded 5. Family members were also removed randomly from the sample.

The IMAGEN cohort (https://www.imagen-project.org/) is a longitudinal study conducted across 8 sites in four European countries, enrolling 2,090 adolescents aged 14 to 22 years. For more information on IMAGEN protocols, see Schumann et al. (2010). This paper analysed three imaging time points (i.e., at ages 14, 19, and 22), from which we included a total sample of 1,825 adolescents (882 males and 942 females). We excluded an MRI time point if the absolute z-score of the neuroimaging metric of interest was over 5. Family members were also removed randomly from the sample.

In both cohorts, we included participants with missing data from the questionnaires used to assess adversity and the neuroimaging data (e.g., participants with only one or two time points). We used probabilistic principal component analysis (PCA) for the adversity scores, and we fit latent growth curve models using Full Information Maximum Likelihood estimation to account for missing data. For the sex analyses, we used sex assigned at birth and excluded the participants who reported being intersex due to the small sample.

### Adversity data

We used the Adverse Adolescent Experiences Framework to conceptualise adversity during adolescence (Pollmann et al., 2025). In our analyses, we included four levels of adversity: intrapersonal, interpersonal with caregivers, interpersonal with peers, and community (Table 1). Details on the items used for each component are provided in Tables 1 and 2 in the Supplementary Material. We selected adversity items that were measured (mainly) at the first time point, reflecting childhood and early adolescence experiences. We defined the four levels of adversity as follows: (1) **Intrapersonal**: Adverse events experienced directly by the adolescent or within the family; (2) **Caregiver**: Perceived lack of warmth and emotional support from caregivers; (3) **Peer**: Experiences of bullying, primarily in school settings; and (4) **Community**: Feelings of unsafety in the neighbourhood.

**Table 1.** Table of the questionnaire details for each component of adversity used in the probabilistic PCA for ABCD and IMAGEN, and the items with the highest variance explained for each component of adversity. [C] is a questionnaire completed by the primary caregiver, and [A] is a questionnaire completed by the adolescent.

|  | ABCD | IMAGEN |
| --- | --- | --- |
| **Intrapersonal adversity** | • KSADS Diagnostic Interview for DSM-5 (Post-Traumatic Stress Disorder) [C]<br>• Victimization Questionnaire [C]<br>• Negative Life Events Questionnaire [A]<br>*Someone in the family arrested.*<br>*One of the caregivers went to jail.*<br>*Trouble with the law for caregiver.*<br>*Family member had drug and/or alcohol problem.*<br>*Family member seriously injured.* | • Life-Events Questionnaire [A]<br>• The Development and Well-Being Assessment Interview [C]<br>*Parental separation.*<br>*Parents divorced.*<br>*Family accident or illness.*<br>*Other stressful life event.*<br>*Serious accident or illness.* |
| **Caregiver adversity** | • PhenX Neighborhood Safety/Crime Survey [A]<br>All items are inverted<br>*Caregiver is able to make me feel better when I am upset.*<br>*Caregiver makes me feel better after talking over my worries with them.*<br>*Caregiver smiles at me very often.*<br>*Caregiver believes in showing their love for me.*<br>*Caregiver is easy to talk to.* | • Childhood Relationships Questionnaire [A]<br>• The Development and Well-Being Assessment Interview [C]<br>*When I was a child I sometimes got the feeling that my mother wished I was never born.*<br>*In childhood I often had the impression that my mother was not listening to me.*<br>*She often tuned me out.*<br>*When I acted bad as a child my mother would, at times, threaten to send me away.* |
|  |  | *In childhood my mother sometimes threatened to leave me or to send me away if I wasn't good.* |
| **Peer adversity** | • KSADS Background Items Survey [A]<br>• Victimization Questionnaire [A]<br>• Negative Life Events Questionnaire [A]<br>Problems with bullying at school or in your neighborhood.<br>Lost a close friend. | • Bully Questionnaire [A]<br>• The Development and Well-Being Assessment Interview [C]<br>*Bullied at school (a student/ peer said or did nasty or unpleasant things to me). Called mean names, was made fun of, or teased in a hurtful way by a student/ peer.* |
| **Community adversity** | • PhenX Neighborhood Safety/Crime Survey [C]<br>• PhenX Neighborhood Safety/Crime Survey [A]<br>All items are inverted<br>*I feel safe walking in my neighborhood, day or night.*<br>*Violence is not a problem in my neighborhood.*<br>*My neighborhood is safe from crime.* | • The Development and Well-Being Assessment Interview [C]<br>*Family stresses: Neighbours or neighbourhood.* |

The conceptualisation of adversity data is currently under debate in the field. First, examining different types of adversity is critical, as lumping diverse experiences into a single adversity score may obscure specific brain-environment relationships (Sheridan & McLaughlin, 2014; Vaidya et al., 2024). Second, while sum scores have long been the norm, following the cumulative risk framework (Evans et al., 2013; McLaughlin et al., 2021), factor analysis approaches are increasingly used to derive latent variables (Moriarity & Slavich, 2025). Notably, McLaughlin et al. (2023) argued that adversity is better conceptualised as a formative construct (i.e., the construct is a result of the items) rather than as a reflective construct (i.e., the construct is the cause of the items), and thus PCA is conceptually more appropriate than CFA/EFA methods. We created adversity scores using probabilistic principal component analysis. Probabilistic PCA reduces data dimensionality while accounting for missing values, enabling more robust estimation of latent variance compared to traditional PCA (Tipping & Bishop, 1999). Probabilistic PCA was applied separately to each adversity component, generating individual-level scores for each. Higher scores reflect higher levels of adversity. Items included in the same component were scaled together (e.g., all items for Intrapersonal adversity). We reported the percentage of variance explained and the main loadings for each component (see Table 3 in the Supplementary Material). We extracted an individual score for each participant for each adversity component.

Questionnaires were completed by both adolescents and primary caregivers; see Table 1 for details (the names of the questionnaires used for each component and cohort are provided in Tables 1 and 2 of the Supplementary Material). The reliability of caregiver reports has been subject to considerable discussion in the literature. Data from several studies revealed discrepancies between adolescent and caregiver reports of violence and trauma exposure (Howard et al., 1999; Oransky et al., 2013; Stover et al., 2010), trauma-related difficulties (Wamser-Nanney & Campbell, 2021), and social victimization experiences (Tang et al., 2025 in ABCD), with caregivers often underestimating their child’s exposure. These studies underscore the need to use multiple sources of information from both the primary caregiver and the adolescents when screening for adverse experiences, as caregivers’ reports were found to better reflect the severity of experiences in some instances (Tang et al., 2025). Thus, when possible, we used items reported by adolescents and caregivers for each component (Table 1). When this combination was not possible or when the items were redundant, we favoured the adolescent report. Finally, we selected items based on their alignment with our conceptual definitions. We attempted to harmonise questions by component across cohorts as much as possible.

Finally, for the Negative Life Events Questionnaire in ABCD (which contributed to both Interpersonal and Peer adversities), we included the valence of the experiences within the scoring for each item. When adolescents rated an event as ‘mostly good’ and as having little to no impact on them, we classified it as the absence of adversity. This distinction is important, as subjective experience has been found to be more strongly associated with risk of psychopathology compared to objective assessment (Danese & Widom, 2020).

### Brain structure data

Grey matter volume and fractional anisotropy were used to measure grey and white matter, respectively, given their widespread use in the literature.

*ABCD.* MRI scans were collected across the 21 research sites using Siemens Prisma, GE 750, and Philips 3T scanners. Scanning protocols were harmonised across sites and scanners. Detailed information on the imaging acquisition protocols and processing methods used is outlined elsewhere (Casey et al., 2018; Hagler et al., 2019).

Grey matter volume was estimated from MRI scans with FreeSurfer (version 7.1.1), and we extracted total cortical grey matter volume from the 68 regions of interest (Desikan et al., 2006). Total cortical grey matter volume was scaled by dividing the metric by 10,000. Fractional anisotropy was measured through diffusion-weighted imaging (DWI). Average fractional anisotropy was extracted from 11 white matter tracts. Fractional anisotropy was scaled by multiplying the metric by 100. For more details on the pipeline used, see Hagler et al., 2019.

*IMAGEN.* MRI scans were collected across the 8 research sites using Siemens Magnetom Verio Syngo MRB17, Philips Achieva, GE DISCOVERY MR750, Siemens Magnetom Trio Tim Syngo MRB17, and Siemens Magnetom Prisma fit scanners. Scanning protocols were harmonised across sites and scanners. Full details of the imaging acquisition protocols and the processing methods used in IMAGEN are outlined elsewhere (Schumann et al., 2010).

Grey matter volume was estimated from MRI scans with FreeSurfer (version 6.0), and we extracted total cortical grey matter volume from the 68 regions of interest (Desikan et al., 2006). Total cortical grey matter volume was scaled by dividing the metric by 10,000. Fractional anisotropy was measured through DWI using the FMRIB Software Library (FSL) (www.fmrib.ox.ac.uk/fsl). The average fractional anisotropy was extracted per participant. Fractional anisotropy was scaled by multiplying the metric by 100.

### Statistical analyses

The goal of this study was to explore the associations between measures of adversity and the trajectories (i.e., intercepts and slopes) of grey and white matter metrics. We aimed to examine the strength of those associations and whether adversity is associated with grey and white matter differently.

#### Growth Mixture Models

Originally, we planned to apply growth mixture models (GMMs) to estimate the trajectories of grey matter volume and fractional anisotropy (Figures 1-4 and Tables 4-7 in the Supplementary Material). GMMs offer an important extension to mixed-effects models by identifying subgroups within the population, each following distinct mean trajectories (Becht & Mills, 2020; Herle et al., 2020). However, our preliminary analyses found no evidence for the existence of subgroups in the trajectories of grey matter volume and fractional anisotropy (i.e., parallel trajectories). As GMMs are known to separate a population into subgroups even when these subgroups do not truly follow distinct trajectories (Bauer, 2007), we decided not to continue with the defined subgroups.

#### Latent growth curve models for grey and white matter metrics

Instead, we modelled their trajectories using latent growth curve models (LGCMs; Stoel et al., 2003; McArdle, 2009). A latent growth curve model is a type of structural equation model that estimates latent variables for the trajectory’s parameters (e.g., intercept and slope). For both grey matter volume and fractional anisotropy, we compared linear, quadratic, and basis models of growth. The basis model estimates individual trajectories by fixing the slope loading at the first and last time points to 0 and 1, respectively, allowing intermediate loadings to be freely determined. Residual variances were constrained to be constant across time points, separately for grey matter volume and fractional anisotropy. We compared the three models’ fit using the likelihood ratio test, including the Akaike Information Criterion (AIC) and the Bayesian Information Criterion (BIC). These parameters penalise models with a higher number of predictors, thereby encouraging the selection of more parsimonious models that explain the data well while avoiding overfitting. We reported AIC_diff_ and BIC_diff_ by subtracting the criterion for the constrained model from the criterion for the free model. Positive values indicate that the preferred model is the free model (i.e., the one with the lower AIC or BIC).

Then, we estimated a bivariate LGCM that included both grey matter volume and fractional anisotropy, allowing the intercepts and slopes to covary. This enabled the observation of the covariance between their trajectories. We used the following guidelines to assess the good fit of the LGCMs: root mean square error of approximation (RMSEA) <0.05 (acceptable, 0.05–0.08), comparative fit index (CFI) >0.97 (acceptable, 0.95–0.97), and standardised root mean square residual (SRMR) <0.05 (acceptable, 0.05–0.10) (Schermelleh-Engel et al., 2003; Mueller & Hancock, 2008).

#### Bivariate latent growth curve models with adversity as predictors

Next, we ran separate bivariate LGCMs for each adversity component, with probabilistic PCA-derived adversity scores predicting the intercepts and slopes of grey matter volume and fractional anisotropy. We evaluated model fit and how each adversity predicted the intercepts and slopes of grey matter volume and fractional anisotropy. Finally, we estimated a comprehensive bivariate LGCM that included the trajectories of grey matter volume and fractional anisotropy, along with the four adversity components (hereafter referred to as the ‘combined model’). The adversity scores were allowed to covary.

In both ABCD and IMAGEN, participants are measured at different sites (21 in ABCD and 8 in IMAGEN) using different scanners. Site correction has become the norm in neuroimaging to disentangle the associations of interest from the site effects (e.g., scanning parameters, participants’ demographics; Bayer et al., 2022). Yet, these many effects render a ‘true’ site correction impossible. In our study, we tried two approaches: (1) include sites as covariates regressed on the intercepts and slopes of both imaging metrics, and (2) run a multigroup model that included sites as groups. The first technique yielded implausible estimates of fractional anisotropy trajectories in both cohorts. Estimates of the mean and variance of the slope of fractional anisotropy indicated no change in the sample, contradicting the plots of the raw data. The multigroup model did not converge for ABCD, likely due to its 21 sites, and we found slightly different results across sites (i.e., groups) for IMAGEN. As the site correction analyses did not yield valid corrections across the two cohorts, we used a Leave One Site Out analysis to assess the robustness of the results by examining the influence of each site (Supplementary Material Figures 7 and 8).

#### Comparison of the regression paths for the grey and white matter metrics

To assess whether adversity measures had differential predictive effects on grey and white matter metrics, we compared a model where structural regression paths (from adversity to brain intercepts and slopes) were freely estimated to a model where the loadings were constrained across brain metrics (e.g., constraining intrapersonal adversity to have equal associations with the intercept of grey matter volume and fractional anisotropy). The raw imaging data were rescaled to constrain the variances of grey matter volume and fractional anisotropy intercepts to 1. This operation enabled a true comparison of the associations between grey matter volume and fractional anisotropy and the different components of adversity. We then tested whether the constrained model had a lower model fit using a likelihood ratio test. If constraining the path coefficients reduced model fit, this would provide evidence that the association between adversity and brain structure differs between grey matter volume and fractional anisotropy.

#### Sex analyses

These analyses were also conducted separately for male and female subsamples. We ran the multigroup bivariate LGCM to determine whether males and females differed in their associations between brain structure and adversity. We also examined the distribution of the four components of adversity, as well as the intercepts and slopes by sex.

All analyses were conducted separately for the ABCD and IMAGEN datasets using R version 4.1.0 (http://www.r-project.org/) and the lavaan package (Rosseel, 2012). All models were fitted using Robust Maximum Likelihood Estimation, with FIML to account for missing data and robust estimation with adjusted standard errors to deal with deviations from normality.

### Data and Code Availability

Data can be requested through [https://nda.nih.gov/] for ABCD and [https://www.imagen-project.org/the-imagen-dataset] for IMAGEN. The code to reproduce the analyses is available on [https://github.com/leacmichel/adversity_grey_white_matter].

## Results

### Characterising adverse experiences in childhood and adolescence

Items measuring adverse experiences were categorised into four components (Intrapersonal, Caregiver, Peer, and Community) through individual probabilistic PCA. The items included in each analysis and the variance explained in each final score are shown in Table 3 in the Supplementary Material. We found that the four components of adversity were not correlated with each other. This result is consistent with the conceptualization of these components as distinct forms of adversity.

The estimation of the Intrapersonal component was the most challenging, as it does not correspond to a single specific form of adversity and, as a result, captured relatively little variance in the data across both cohorts (9% in ABCD and 15% in IMAGEN). Given the weak pattern of correlations among the items, the Intrapersonal adversity component expresses the items with the highest loadings. In ABCD, this component primarily reflected traumatic experiences such as injuries, substance use issues in the family, and problems with the law in the family. In IMAGEN, it was more reflective of questions related to parental separation, divorce, accidents, and illness in the family.

The Caregiver component explained a substantial proportion of variance in the data in both cohorts. In ABCD, it was moderately loaded (36%) on all the items measuring caregivers’ warmth. In IMAGEN, it was moderately loaded (34%) on questions related to the perception of attachment to their caregivers.

The Peer component explained 27% of the variance in the ABCD data and 50% in the IMAGEN data. This is explained by the fact that some questions in ABCD had more than 80% missing values, thereby reducing the explained variance. In both cohorts, this component primarily captured school bullying. In ABCD, the question related to loss of friendships also showed a high loading on this component.

The Community component focused on feelings of safety in the neighbourhood. In ABCD, this component explained 64% of the variance in the data but centred on caregivers’ perceptions. In IMAGEN, the component only included a single item about family stress due to the neighbourhood.

### Trajectories of brain development

Trajectories of brain development captured using latent growth curve analyses are shown in Figure 1. The basis model was preferred over the linear and quadratic models for total grey matter volume in both ABCD (Δχ²(1) = 466.21, p < 0.001; AIC_diff_ = 594; BIC_diff_ = 587 in favour of the basis model) and IMAGEN (Δχ²(1) = 212, p < 0.001; AIC_diff_ = 366; BIC_diff_ = 362 in favour of the basis model) and for fractional anisotropy in IMAGEN (Δχ²(1) = 49.57, p < 0.001; AIC_diff_ = 103; BIC_diff_ = 97 in favour of the basis model), while the linear model was preferred for the average fractional anisotropy trajectories in ABCD. This indicates that changes in grey matter volume and fractional anisotropy (only in IMAGEN) are not linear.

**Figure 1.**
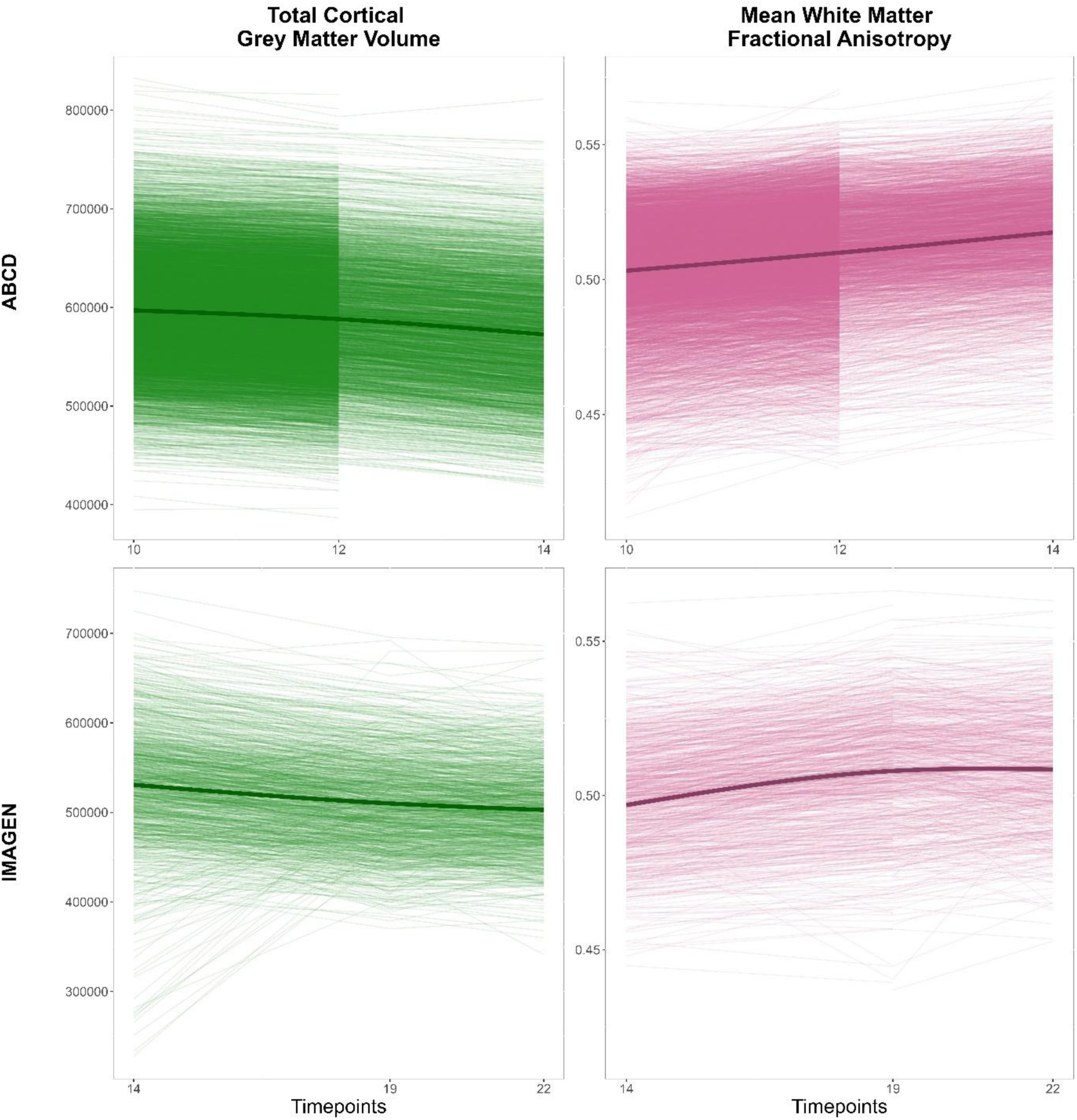
Trajectories of total cortical grey matter volume and mean fractional anisotropy in ABCD and IMAGEN. The darker lines represent the population trajectory approximated with a generalised additive model using three splines. The x-axis represents the three time points and the corresponding average ages.

The bivariate latent growth curve model, including total grey matter volume and average fractional anisotropy, fitted the data well in both ABCD and IMAGEN (Tables 2 and 3). In ABCD and IMAGEN, the variance of the slope for grey matter volume was negative in some of the bivariate latent growth curve models. To address this, we constrained the slope variance of grey matter to be positive. Interpretations for negative slopes in grey matter volume will be provided throughout, as only 2 participants (0.02% of the sample) showed a positive slope in ABCD and 153 (8%) in IMAGEN. For fractional anisotropy, interpretations will be provided throughout for positive slopes, as no participants showed a negative slope in ABCD and only 2 participants (0.1%) in IMAGEN did so.

**Table 2.**
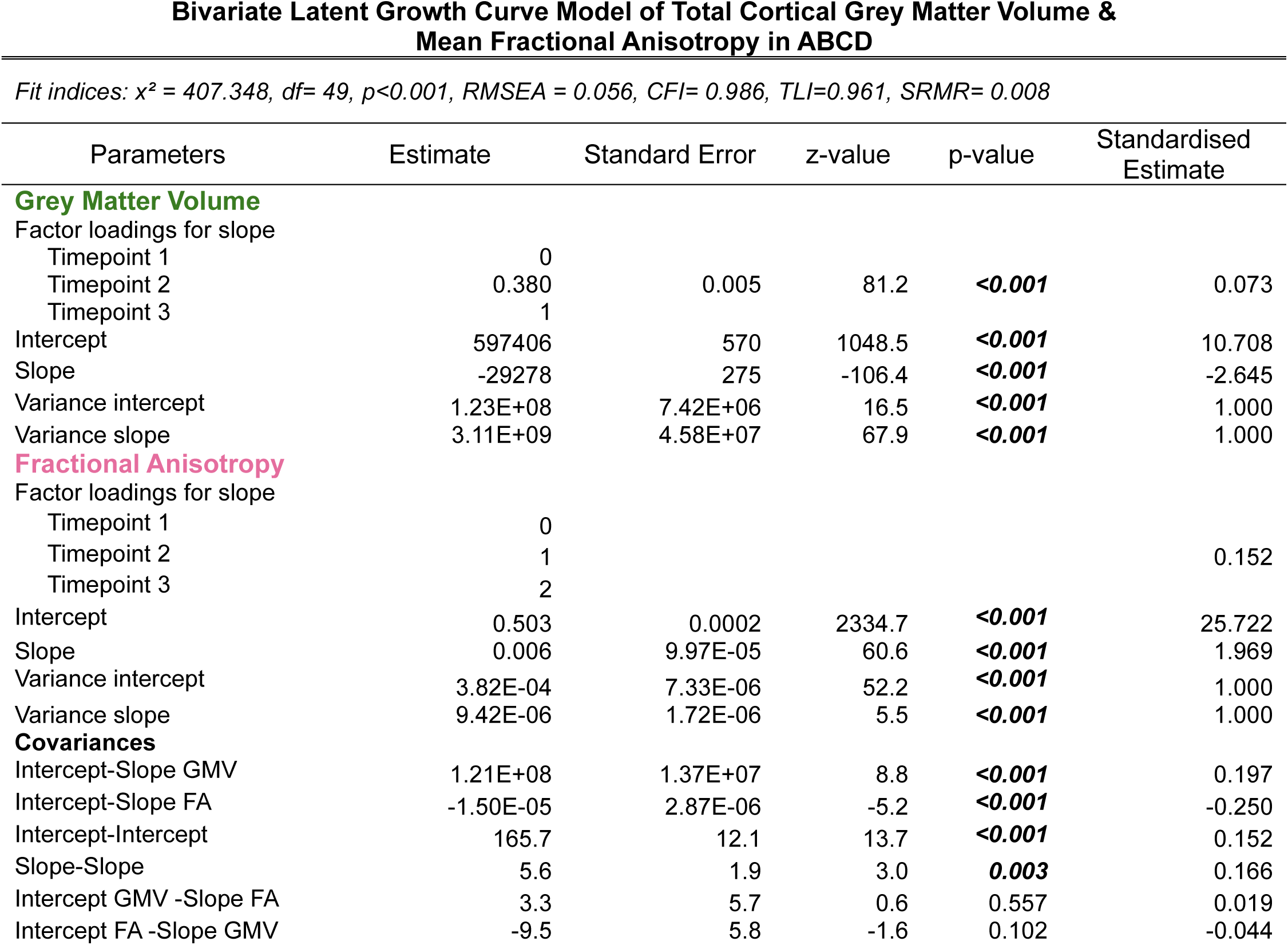
Parameter estimates of the bivariate latent growth curve model of total cortical grey matter volume and mean fractional anisotropy in ABCD.

| Parameters | Estimate | Standard Error | z-value | p-value | Standardised Estimate |
| --- | --- | --- | --- | --- | --- |
| **Grey Matter Volume** |  |  |  |  |  |
| Factor loadings for slope |  |  |  |  |  |
| Timepoint 1 | 0 |  |  |  |  |
| Timepoint 2 | 0.380 | 0.005 | 81.2 | **<0.001** | 0.073 |
| Timepoint 3 | 1 |  |  |  |  |
| Intercept | 597406 | 570 | 1048.5 | **<0.001** | 10.708 |
| Slope | -29278 | 275 | -106.4 | **<0.001** | -2.645 |
| Variance intercept | 1.23E+08 | 7.42E+06 | 16.5 | **<0.001** | 1.000 |
| Variance slope | 3.11E+09 | 4.58E+07 | 67.9 | **<0.001** | 1.000 |
| **Fractional Anisotropy** |  |  |  |  |  |
| Factor loadings for slope |  |  |  |  |  |
| Timepoint 1 | 0 |  |  |  |  |
| Timepoint 2 | 1 |  |  |  | 0.152 |
| Timepoint 3 | 2 |  |  |  |  |
| Intercept | 0.503 | 0.0002 | 2334.7 | **<0.001** | 25.722 |
| Slope | 0.006 | 9.97E-05 | 60.6 | **<0.001** | 1.969 |
| Variance intercept | 3.82E-04 | 7.33E-06 | 52.2 | **<0.001** | 1.000 |
| Variance slope | 9.42E-06 | 1.72E-06 | 5.5 | **<0.001** | 1.000 |
| **Covariances** |  |  |  |  |  |
| Intercept-Slope GMV | 1.21E+08 | 1.37E+07 | 8.8 | **<0.001** | 0.197 |
| Intercept-Slope FA | -1.50E-05 | 2.87E-06 | -5.2 | **<0.001** | -0.250 |
| Intercept-Intercept | 165.7 | 12.1 | 13.7 | **<0.001** | 0.152 |
| Slope-Slope | 5.6 | 1.9 | 3.0 | **0.003** | 0.166 |
| Intercept GMV -Slope FA | 3.3 | 5.7 | 0.6 | 0.557 | 0.019 |
| Intercept FA -Slope GMV | -9.5 | 5.8 | -1.6 | 0.102 | -0.044 |

**Table 3.**
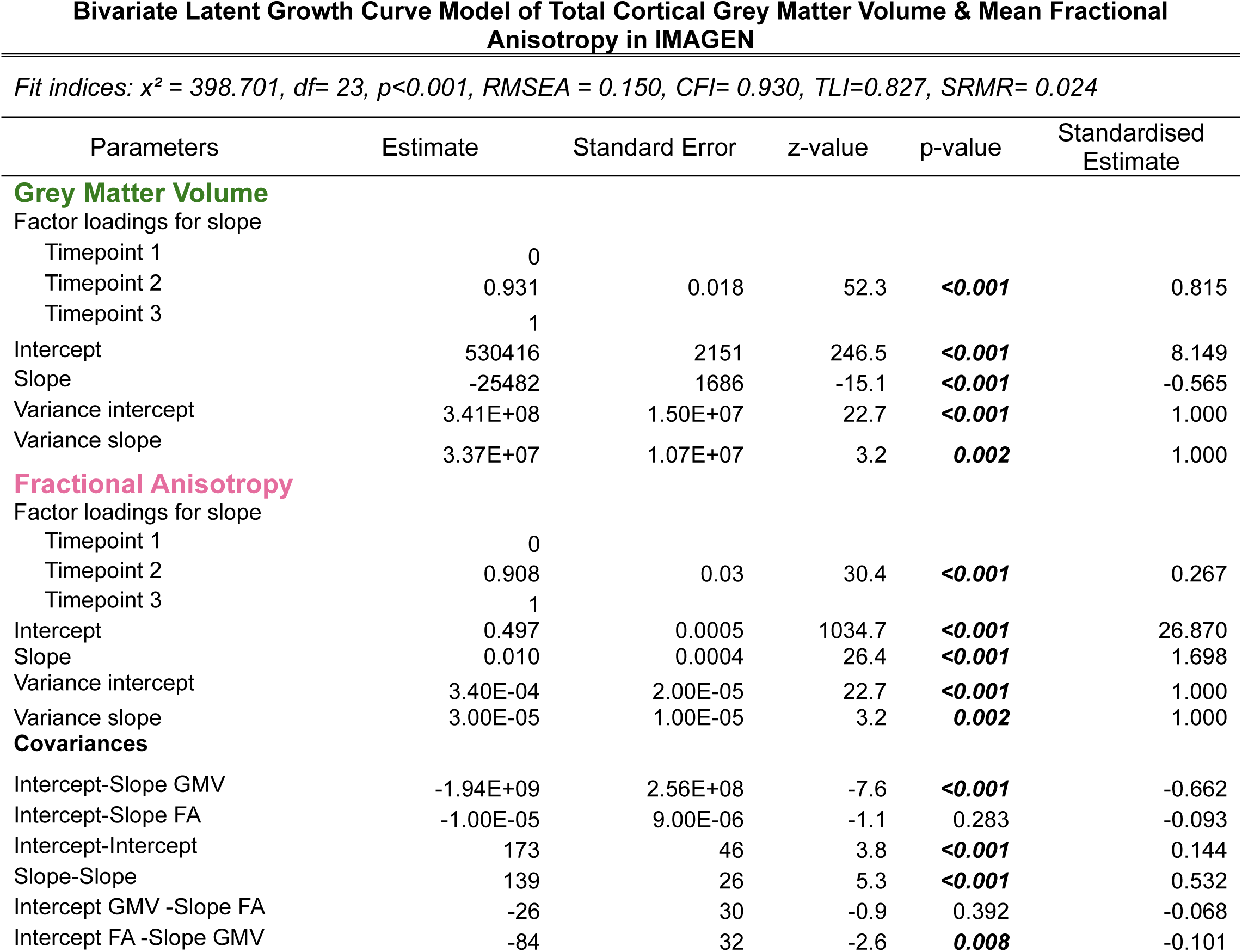
Parameter estimates of the bivariate latent growth curve model of total cortical grey matter volume and mean fractional anisotropy in IMAGEN.

The model allowed the intercepts and slopes of both metrics to covary. The development of grey matter volume was found to correlate with the development of fractional anisotropy, via intercept-intercept, slope-slope, and, to a lesser extent, intercept-slope correlations.

### Adversity predicted the intercepts of grey matter volume and fractional anisotropy, but not the slopes

All individual models (i.e., a model with one adversity component predicting both grey matter volume and fractional anisotropy) and the combined model (i.e., a model with the four adversity components predicting both grey matter volume and fractional anisotropy) fitted the data well in ABCD and IMAGEN. The results of the individual models matched those of the combined model; thus, we focus on the combined model’s results here (see Tables 8-15 in the Supplementary Material for the summary parameters of the individual models).

In both cohorts, only Caregiver, Peer, and Community adversities predicted the intercepts of grey and white matter. Intrapersonal adversity did not predict intercepts or slopes in either metric. Surprisingly, adversity predicted only the intercepts, not the slopes, for both grey matter volume and fractional anisotropy in both cohorts.

In ABCD, adversity predicted the intercepts of both grey matter volume and fractional anisotropy (Figure 2 in the paper and Figure 5 in the Supplementary Material). Both the Peer and Community components negatively predicted the intercepts for total grey matter volume (-0.039 and -0.104, respectively), suggesting that adolescents who experienced bullying or felt unsafe in their neighbourhood during childhood showed lower total grey matter volume at age 10. For fractional anisotropy, both Caregiver (-0.031) and Community (-0.064) adversities negatively predicted the intercepts of fractional anisotropy. This finding indicates that adolescents who perceived less warmth in their relationships with their caregivers or who felt unsafe in their neighbourhoods showed lower fractional anisotropy at age 10. These results remained robust in the Leave-One-Site-Out analysis across the four associations (Figure 7 in the Supplementary Material).

**Figure 2.**
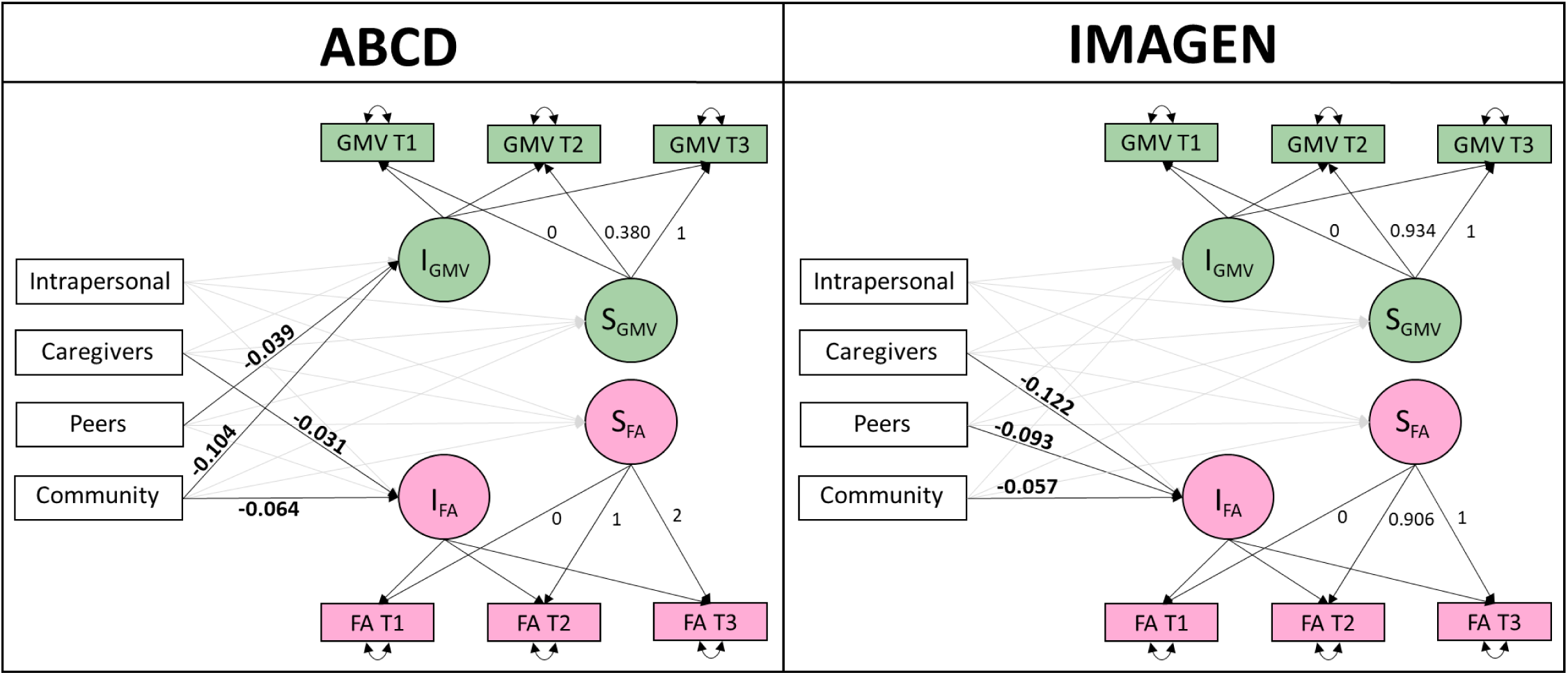
Path diagrams of the bivariate latent growth curve model for grey matter volume and fractional anisotropy with the four components of adversity as predictors in both ABCD and IMAGEN. We displayed the path coefficients for the significant associations (p<0.05) between adversity and grey and white matter development in bold. Detailed statistics for each model are presented in Tables 4 & 5.

**Table 4.** Parameter estimates of the bivariate latent growth curve model with the four levels of adversity predicting the intercepts and slopes of total cortical grey matter volume and mean fractional anisotropy in ABCD.

| Combined model with adversity (ABCD) |  |  |  |  |  |  |  |
| --- | --- | --- | --- | --- | --- | --- | --- |
| *Fit indices: $\chi^2 = 131.829$, $df = 18$, $p < 0.001$, $RMSEA = 0.047$, $CFI = 0.996$, $TLI = 0.991$, $SRMR = 0.010$* |  |  |  |  |  |  |  |
| Parameters | Estimate | Std.error | z-value | p-value | Std.estimate | CI lower | CI upper |
| **Intrapersonal** |  |  |  |  |  |  |  |
| Intercept GMV | -645 | 497 | -1.30 | 0.194 | -0.014 | -1620 | 329 |
| Intercept FA | -7.95E-05 | 1.91E-04 | -0.42 | 0.678 | -0.005 | -4.54E-04 | 2.95E-04 |
| Slope GMV | 149 | 230 | 0.65 | 0.517 | 0.016 | -301 | 600 |
| Slope FA | -2.35E-05 | 7.55E-05 | -0.31 | 0.756 | -0.009 | -1.71E-04 | 1.25E-04 |
| **Caregiver** |  |  |  |  |  |  |  |
| Intercept GMV | -751 | 516 | -1.46 | 0.145 | -0.016 | -1762 | 260 |
| Intercept FA | -5.03E-04 | 1.83E-04 | -2.74 | **0.006** | -0.031 | -8.62E-04 | -1.44E-04 |
| Slope GMV | 351 | 232 | 1.51 | 0.130 | 0.038 | -103 | 805 |
| Slope FA | 3.09E-05 | 8.83E-05 | 0.35 | 0.726 | 0.012 | -1.42E-04 | 2.04E-04 |
| **Peer** |  |  |  |  |  |  |  |
| Intercept GMV | -3799 | 1040 | -3.65 | **<0.001** | -0.039 | -5836 | -1761 |
| Intercept FA | -9.68E-05 | 3.83E-04 | -0.25 | 0.800 | -0.003 | -8.46E-04 | 6.53E-04 |
| Slope GMV | 53 | 513 | 0.10 | 0.917 | 0.003 | -952 | 1058 |
| Slope FA | 8.90E-05 | 1.87E-04 | 0.47 | 0.635 | 0.016 | -2.78E-04 | 4.56E-04 |
| **Community** |  |  |  |  |  |  |  |
| Intercept GMV | -5980 | 584 | -10.25 | **<0.001** | -0.104 | -7124 | -4836 |
| Intercept FA | -1.29E-03 | 2.14E-04 | -6.05 | **<0.001** | -0.064 | -1.71E-03 | -8.73E-04 |
| Slope GMV | -88 | 289 | -0.30 | 0.762 | -0.008 | -655 | 480 |
| Slope FA | 1.90E-04 | 1.05E-04 | 1.80 | 0.071 | 0.059 | -1.63E-05 | 3.97E-04 |

**Table 5.** Parameter estimates of the bivariate latent growth curve model with the four levels of adversity predicting the intercepts and slopes of total cortical grey matter volume and mean fractional anisotropy in IMAGEN.

| Parameters | Estimate | Std.error | z-value | p-value | Std.estimate | CI lower | CI upper |
| --- | --- | --- | --- | --- | --- | --- | --- |
| **Intrapersonal** |  |  |  |  |  |  |  |
| Intercept GMV | -2091 | 1974 | -1.059 | 0.290 | -0.040 | -5961 | 1778 |
| Intercept FA | 0.0005 | 0.0004 | 1.352 | 0.176 | 0.035 | -0.0002 | 0.0013 |
| Slope GMV | 2822 | 1650 | 1.710 | 0.087 | 0.077 | -412 | 6056 |
| Slope FA | 0.0002 | 0.0003 | 0.700 | 0.484 | 0.044 | -0.0004 | 0.0008 |
| **Caregiver** |  |  |  |  |  |  |  |
| Intercept GMV | -508 | 937 | -0.542 | 0.588 | -0.016 | -2345 | 1329 |
| Intercept FA | -0.0011 | 0.0003 | -4.097 | **<0.001** | -0.122 | -0.0017 | -0.0006 |
| Slope GMV | 629 | 625 | 1.005 | 0.315 | 0.028 | -597 | 1854 |
| Slope FA | 0.0001 | 0.0002 | 0.685 | 0.493 | 0.038 | -0.0002 | 0.0004 |
| **Peer** |  |  |  |  |  |  |  |
| Intercept GMV | -116 | 1448 | -0.080 | 0.936 | -0.003 | -2954 | 2722 |
| Intercept FA | -0.0011 | 0.0003 | -3.739 | **<0.001** | -0.093 | -0.0017 | -0.0005 |
| Slope GMV | 620 | 985 | 0.629 | 0.529 | 0.021 | -1311 | 2550 |
| Slope FA | 0.0004 | 0.0003 | 1.621 | 0.105 | 0.113 | -0.0001 | 0.0010 |
| **Community** |  |  |  |  |  |  |  |
| Intercept GMV | 2223 | 6045 | 0.368 | 0.713 | 0.009 | -9626 | 14071 |
| Intercept FA | -0.0041 | 0.0017 | -2.439 | **0.015** | -0.057 | -0.0074 | -0.0008 |
| Slope GMV | -3518 | 4136 | -0.851 | 0.395 | -0.020 | -11625 | 4589 |
| Slope FA | 0.0009 | 0.0012 | 0.781 | 0.435 | 0.040 | -0.0014 | 0.0032 |

In IMAGEN, only fractional anisotropy was predicted by adversity (Figure 2 in the paper and Figure 6 in the Supplementary Material). Caregiver component (-0.122), Peer component (-0.093), and Community component (-0.057) were all associated with lower fractional anisotropy at age 14. These results suggest that adolescents who experienced higher adversity in childhood in these three domains showed lower fractional anisotropy at 14. These results remained robust in the Leave One Site Out analysis for the associations with Caregiver and Peer adversity. For Community adversity, the negative association was not observed when data from Dresden or Paris were removed (Figure 8 in the Supplementary Material).

Thus, in both samples, we found that Caregiver and Community adversities during childhood were associated with lower fractional anisotropy. In ABCD, Community adversity was also associated with lower grey matter volume. Peer adversity was linked with lower grey matter volume in ABCD and lower fractional anisotropy in IMAGEN.

### Adversity predicts grey and white matter differently

To test whether adversity predicted grey and white matter trajectories similarly, we constrained the path coefficients to be the same across grey matter volume and fractional anisotropy. In both cohorts, the unconstrained (free) model was preferred over the fully constrained model (ABCD: Δχ²(8) = 22.27, p = 0.004; AIC_diff_ = 7; BIC_diff_ = -51; IMAGEN: Δχ²(8) = 26.69, p < 0.001; AIC_diff_ = 9; BIC_diff_ = -35), suggesting that adversity is associated with grey and white matter differently.

To determine whether these differential associations were specific to particular forms of adversity, we conducted follow-up analyses by constraining the model one adversity component at a time. In ABCD, the free model was preferred for both Peer (Δχ²(2) = 6.39, p = 0.04; AIC_diff_ = 3; BIC_diff_ = -12) and Community adversities (Δχ²(2) = 12.51, p = 0.002; AIC_diff_ = 9; BIC_diff_ = -6), suggesting that these adversities were differently associated with the trajectories of grey matter volume and fractional anisotropy, with a stronger association observed for grey matter volume. In IMAGEN, the free model was preferred only for Caregiver adversity (Δχ²(2) = 13.21, p = 0.001; AIC_diff_ = 8; BIC_diff_ = -3) compared to the models that constrained the path coefficients for the intercepts and slopes. This suggests that Caregiver adversity has unique associations on grey and white matter, with a stronger association for fractional anisotropy. We also explored a model where only the path coefficients to the intercepts were constrained, since we found significant associations only between adversity and the intercepts, not the slopes. We found that the free model was also preferred for the Peer (Δχ²(1) = 5.15, p = 0.02; AIC_diff_ =2; BIC_diff_ =-3) and Community (Δχ²(1) = 4.16, p = 0.04; AIC_diff_ =0; BIC_diff_ =-5) components in IMAGEN. Both adversities also showed a stronger association with fractional anisotropy intercepts. Figure 3 illustrates that higher scores for adversity in these components were associated with grey matter volume and fractional anisotropy trajectories in ABCD and IMAGEN.

**Figure 3.**
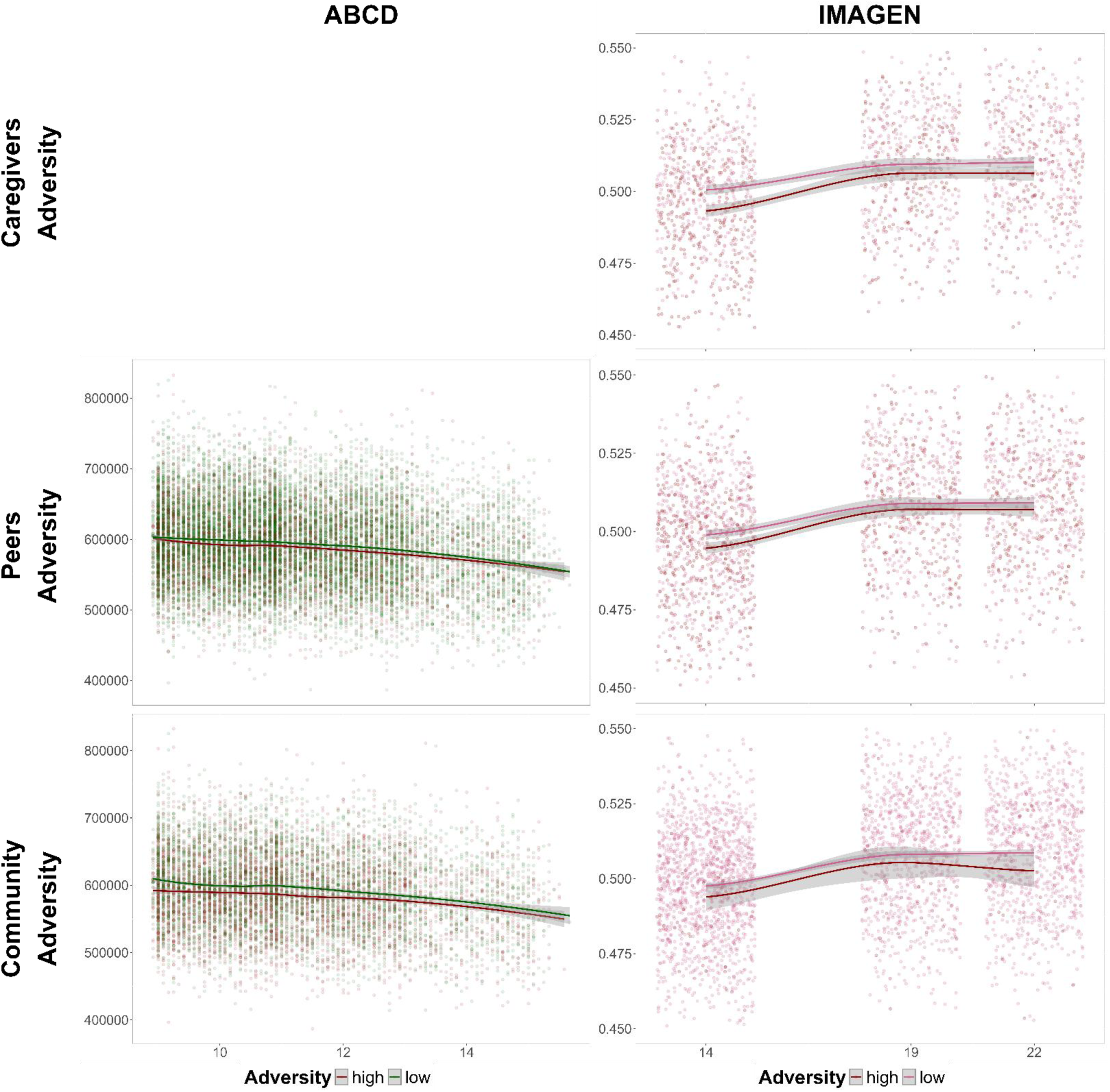
Trajectories of total grey matter volume (in green) and average fractional anisotropy (in pink) across age (x-axis) for a group with high adversity (25% highest score – red line) and a group with low adversity (25% lower score) in both ABCD and IMAGEN. We displayed the three components of adversity that showed differences in their associations with grey matter volume and fractional anisotropy. For each component in each cohort, we show the metric that was significantly associated with the specific adversity. Each trajectory is represented through a generalised additive model using three splines. In IMAGEN, the datapoints have been represented with some degree of jitter.

The differences in model fit criteria diverge in their conclusions across all likelihood-ratio tests, with positive AIC differences and negative BIC differences. AIC favours more complex models while BIC favours more parsimonious models, as BIC penalises additional parameters (here the free model) more heavily compared to the AIC. This discordance suggests small differences in effect sizes. For instance, in ABCD, Peer significantly predicts the grey matter volume intercepts (-0.039), but not the fractional anisotropy intercepts (-0.003). In the constrained model, these associations are both constrained to be equal, so that Peer significantly predicts both the intercepts of grey matter volume and fractional anisotropy (-0.022).

Finally, we assessed the strength of the associations between adversity and the trajectories of grey and white matter. In ABCD, adversity explained three times more variance (R-squared) in the intercepts (R^2^ = 0.013) but not the slopes (R^2^ = 0.002) of total grey matter volume compared to the intercepts (R^2^ = 0.005) and slopes (R^2^ = 0.004) of average fractional anisotropy. Conversely, in IMAGEN, adversity explained more variance in the intercepts (R^2^ = 0.031) and slopes (R^2^ = 0.020) of average fractional anisotropy compared to the intercepts (R^2^ = 0.002) and slopes (R^2^ = 0.008) of total grey matter volume. These findings suggest that childhood adversity is more strongly associated with grey matter volume in ABCD but with fractional anisotropy in IMAGEN. These differences could result from variations in ages, items, or population. We explore this further in the Discussion.

### Adversity predicts grey and white matter trajectories differently in males and females

We found small-to-moderate differences in the distribution of adversity between males and females across the four components, particularly in IMAGEN (see Figures 9 and 11 in the Supplementary Material). We also observed differences in the probabilistic PCA loadings between males and females. For instance, the Intrapersonal component in IMAGEN appears to be driven by slightly different items in males and in females (see Figures 10 and 12 in the Supplementary Material). Thus, we decided to use the probabilistic PCA scores from the full sample to ensure that observed differences were not driven by sex-specific probabilistic PCA scores (i.e., sex differences in the frequency and the way participants experienced adversity).

The combined models, stratified by sex, revealed distinct patterns in both ABCD and IMAGEN (Figure 4 in the paper and Tables 16 and 17 in the Supplementary Material, respectively). In ABCD, the sex-specific analyses largely replicated the population-level results for Peer and Community adversities, suggesting that these associations are robust across sex. However, we found an association between the intercepts of fractional anisotropy and Caregiver adversity in females, but not in males. For Caregiver adversity, we also found that females showed a negative association between Caregiver adversity and the grey matter volume intercepts (-0.041), whereas males showed a positive association between Caregiver adversity and the grey matter volume slopes (0.121). This suggests that male adolescents who experienced higher levels of Caregiver adversity in childhood had a slower decrease in grey matter volume between ages 10 and 14, whereas females who experienced greater Caregiver adversity had lower total volume of grey matter at 10. Finally, in males, we found that Community adversity was positively associated with the slope of fractional anisotropy (0.119), indicating that male adolescents who felt more unsafe in their neighbourhood experienced faster increases in fractional anisotropy during adolescence.

**Figure 4.**
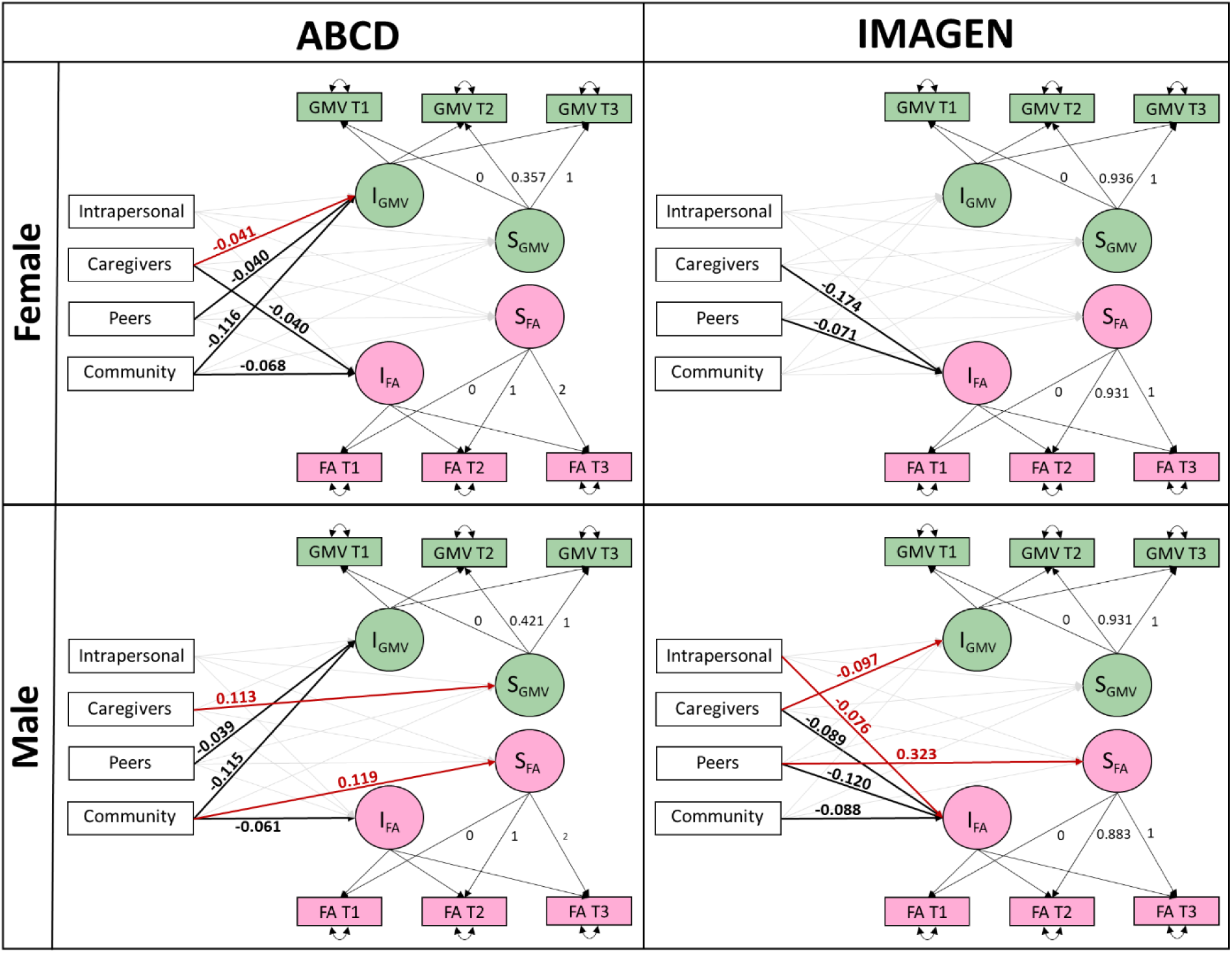
Path diagrams of the bivariate latent growth curve model for grey matter volume and fractional anisotropy with the four components of adversity as predictors in both ABCD and IMAGEN by sex. We displayed the path coefficients for the significant associations (p<0.05) between adversity and grey and white matter development. The black paths are those we found significant in both males and females in bold. The red paths are those significant only in either females or males.

In IMAGEN, the sex-stratified analysis replicated the negative association between Caregiver and Peer adversities and the intercepts of fractional anisotropy. However, the analysis also yielded results that diverged from the population-level model. Intrapersonal adversity negatively predicted the intercepts of fractional anisotropy in males (-0.076). The negative association indicates that males experiencing higher levels of Intrapersonal adversity (e.g., parental separation, accident/illness in the family) have smaller total cortical grey matter volume at 14. Caregiver adversity negatively predicted fractional anisotropy in both sexes, with a stronger association in females (-0.174) than in males (-0.089) and negatively predicted the intercepts of grey matter volume in males (-0.097). This suggests that adolescents who felt less affection from their caregivers showed lower levels of fractional anisotropy; in males, this experience is also associated with lower grey matter volume. Peer adversity negatively predicted the intercepts of fractional anisotropy (-0.120) in males and (-0.071) in females, while also positively predicting its slopes (0.323) in the male subsample. These findings indicate that adolescents who experienced more bullying in their childhood showed lower fractional anisotropy at age 14, followed by a steeper subsequent increase up to age 22. Community adversity negatively predicted the intercepts of fractional anisotropy (-0.088) only in the male sample.

We also reproduced the analyses, constraining the path coefficients to be the same across grey matter volume and fractional anisotropy. We wanted to test whether adversity predicted grey matter volume and fractional anisotropy trajectories similarly in the male and female subsamples. In ABCD, for the male subsample, the unconstrained models were preferred over the constrained models for the combined model (i.e., constraining all four components of adversity, Δχ²(8) = 25.02, p = 0.002; AIC_diff_ = 9; BIC_diff_ = -48), as well as for the model constraining Community adversity (Δχ²(2) = 14.47, p < 0.001; AIC_diff_ = 11; BIC_diff_ = -3). For females, the free model was preferred for Community adversity (Δχ²(2) = 6.63, p = 0.04; AIC_diff_ = 2; BIC_diff_ = -12). When exploring models that constrained only the intercepts, the unconstrained model was preferred for Peer adversity (Δχ²(1) =3.93, p = 0.048; AIC_diff_ =2; BIC_diff_ =-5) in the male subsample as well. Thus, the stronger associations between Community adversity and grey matter volume were confirmed in the sex-stratified analyses, but not fully for those with Peer adversity, despite replicating its negative association with grey matter volume intercepts.

In IMAGEN, in the male subsample, the unconstrained models were preferred only for the models constraining Peer adversity (Δχ²(2) = 6.33, p = 0.042; AIC_diff_ =2; BIC_diff_ =-8). In contrast, in the female subsample, the unconstrained model was favoured for the combined model (Δχ²(8) = 18.94, p = 0.015; AIC_diff_ =0; BIC_diff_ =-44) and the model constraining Caregiver adversity (Δχ²(2) = 11.77, p = 0.003; AIC_diff_ =6; BIC_diff_ =-5). The sex-stratified analysis only (partially) reproduces the full sample result for Caregiver and Peer, as Caregiver adversity was more strongly associated with fractional anisotropy in females but not in males, and Peer adversity was more strongly associated with fractional anisotropy in males but not in females.

### Summary of associations found between brain structure and adversity by sex

Summary of associations found between brain structure and adversity by sex.
| ABCD | Intercepts/Slopes Grey Matter |  |  |  |  |  | Intercepts/Slopes White Matter |  |  |  |  |  |
| --- | --- | --- | --- | --- | --- | --- | --- | --- | --- | --- | --- | --- |
|  | General |  | Female |  | Male |  | General |  | Female |  | Male |  |
| Intrapersonal | - | - | - | - | - | - | - | - | - | - | - | - |
| Caregivers | - | - | ✓ | - | - | ✓ | ✓ | - | ✓ | - | - | - |
| Peers | ✓ | - | ✓ | - | ✓ | - | - | - | - | - | - | - |
| Community | ✓ | - | ✓ | - | ✓ | - | ✓ | - | ✓ | - | ✓ | ✓ |

| IMAGEN | Intercepts/Slopes Grey Matter |  |  |  |  |  | Intercepts/Slopes White Matter |  |  |  |  |  |
| --- | --- | --- | --- | --- | --- | --- | --- | --- | --- | --- | --- | --- |
|  | General |  | Female |  | Male |  | General |  | Female |  | Male |  |
| Intrapersonal | - | - | - | - | - | - | - | - | - | - | ✓ | - |
| Caregivers | - | - | - | - | ✓ | - | ✓ | - | ✓ | - | ✓ | - |
| Peers | - | - | - | - | - | - | ✓ | - | ✓ | - | ✓ | ✓ |
| Community | - | - | - | - | - | - | ✓ | - | - | - | ✓ | - |

## Discussion

This study explored the associations between adversity and structural metrics of grey and white matter in adolescence in the ABCD (N=9,994 at ages 10, 12, and 14) and IMAGEN (N=2,090 at ages 14, 19, and 22) studies. In both ABCD and IMAGEN, childhood adversity showed distinct associations with grey and white matter structures in adolescence. In the younger ABCD sample, adversities associated with the Peer (e.g., bullying) and Community (e.g., feeling unsafe in the neighbourhood) components were more strongly associated with grey matter volume than with fractional anisotropy. Adolescents who experienced more bullying in their childhood, or who reported feeling less safe in their neighbourhood, showed lower levels of grey matter volume at age 10. In contrast, in the older IMAGEN sample, Caregiver, Peers and Community adversities were significantly differentially associated with the intercepts of grey matter volume and fractional anisotropy in adolescence. For instance, adolescents who reported feeling unwanted by their caregivers during childhood had lower fractional anisotropy at age 14. Interestingly, no clear pattern favouring either grey or white matter as more sensitive to adversity emerged. In ABCD, adversity explained more variance in grey matter volume, whereas in IMAGEN, it explained more variance in fractional anisotropy. This suggests that while adversity is associated with both grey and white matter structures, the type of adversity and timing of exposure may determine which neural systems are most affected, although systematic differences between the two samples may also contribute to these divergent patterns.

In our results, the four components of adversity were associated only with the intercepts of the brain trajectories, and not with their slopes (i.e., interindividual rates of change) in either cohort. While the absence of associations with slopes may seem surprising, associations with intercepts may reflect the cumulative effects of earlier experiences and prior brain changes. Therefore, the lack of detectable slope effects in adolescence should not be taken as evidence that adversity does not predict brain development. Instead, it is more likely that most changes in brain structure occurred prior to adolescence, resulting in stronger effects on intercepts. In the literature, studies have reported associations between early adversity and longitudinal changes (i.e., slopes) in subcortical and, to some extent, in grey and white matter structures (Beck et al., 2025; Breslin et al., 2024; Rakesh et al., 2023; Sheridan et al., 2022; Thijssen et al., 2024). These discrepancies with our findings could result from the samples, the conceptualisation of adversity, or the analyses.

Alternatively, associations between adversity and slopes may be better observed by separating males and females. We observed sex-specific associations between adversity and grey and white matter structures in females and males, despite using population scores to achieve a similar distribution of adversity across sexes. Contrary to what we found in the main sample, sex-stratified analyses displayed associations between the components of adversity and the slopes. For instance, in IMAGEN, male adolescents who reported higher levels of Peer adversity showed a greater increase in fractional anisotropy. These findings align with prior studies showing the importance of sex-stratified analyses when examining the associations of adversity with grey matter (Everaerd et al., 2012; Kelly et al., 2015; Peng et al., 2025) and white matter metrics (Teicher et al., 2004). One potential explanation for the observed sex differences and the associations with slopes is the different brain maturational timelines between females and males (Kaczkurkin et al., 2019). There is some evidence that females show an earlier development in grey matter (Carrick et al., 2025; Fuhrmann et al., 2022; Lenroot et al., 2007) and white matter maturation (Asato et al., 2010; Carrick et al., 2025), potentially altering the windows of neurobiological sensitivity to adversity. However, the limited number of longitudinal studies explicitly examining sex differences in brain structure trajectories limits our interpretation of the findings. In addition, with our models, we cannot determine whether these sex differences stem from differences in brain structure or in how adversity is associated with brain structure. Further work is needed to disentangle the roots of those sex differences (see Bath, 2020, for a review on the underlying factors and implications of such questions).

Studies focusing on the effect of adversity on brain development often use the Stress Acceleration hypothesis to interpret their findings, which hypothesises that experiences of adversity, particularly threat, accelerate the pace of development (Callaghan & Tottenham, 2016; Colich et al., 2020). Such accelerated maturation may be an adaptation to adversity in the short term but might increase vulnerability to later mental health problems. In line with this account, Beck et al. (2025) found that different types of adversity were associated with slower or faster apparent brain ageing (i.e., brain age gap) in the ABCD cohort. Emotional neglect (∼ Caregiver adversity) was associated with smaller brain age gaps (i.e., slower development), whereas trauma exposure (∼ Intrapersonal adversity) and neighbourhood safety (∼ Community adversity) were associated with larger brain age gaps (i.e., accelerated development). This pattern was partially observed in our study. In ABCD, adolescents who experienced higher levels of Community adversity (i.e., feeling unsafe) showed, on average, lower grey matter volume at baseline, which may indicate that they were already further along the trajectory of grey matter reduction (i.e., a faster prior rate of reduction). Likewise, we found that adolescents who reported greater Caregiver adversity (i.e., emotional neglect) showed, on average, lower fractional anisotropy at baseline, which may reflect faster development as fractional anisotropy typically increases during this period. Longer longitudinal samples, more heterogeneous populations and assessments over age ranges best aligned with the periods of most rapid neural change are needed to provide greater certainty.

For our study, we conceptualised adversity using the Adolescent Adversity Experiences Framework (Pollmann et al., 2025). While this approach diverges from other models like ACEs (Adverse Childhood Experiences; Felitti et al., 1998) or the threat and deprivation framework (Sheridan & McLaughlin, 2014), our multilevel structure highlights the multidimensional nature of adversity and allows for the possibility that different types of adversity relate to neurodevelopment through distinct mechanisms. Consistent with this idea, we found that Peer and Community adversities were more strongly associated with grey matter volume (in ABCD), whereas Caregiver adversity was more strongly associated with fractional anisotropy (in IMAGEN). Yet, pinpointing the mechanisms underlying these distinct associations remains challenging in human studies. Animal studies show that adversity impacts multiple neuronal mechanisms, which would be reflected through both grey and white matter metrics in human studies. For instance, Lehmann et al. (2017) reported that mice exposed to chronic social defeat (∼ Peer adversity) showed decreased myelination in their prefrontal cortex (∼ lower fractional anisotropy), alongside changes in gene expression related to neurite degeneration, neuroglial cell death, and decreased cell proliferation (∼ lower grey matter volume). Future research should incorporate specific measures of grey and white matter microstructure that better capture specific cellular components, such as Soma And Neurite Density Imaging (SANDI; Palombo et al., 2020), to provide a more comprehensive picture of the neurobiological mechanisms associated with different types of adversity.

A major strength of this study is its use of two large-scale, longitudinal cohorts spanning the beginning (ABCD) and end of adolescence (IMAGEN). However, while we observed differences in the associations between adversity and grey and white matter structures across the two cohorts, several factors limit the direct comparison of the results. These differences in findings could be due to at least three explanations: (1) ABCD and IMAGEN differ in demographic, cultural characteristics, and scanner types; (2) the cohorts were scanned at different developmental windows (ABCD: 10-12-14 years; IMAGEN: 14-19-22 years); and (3) measures of adversity were reported through different questionnaires that might capture slightly different experiences. Future research will need to disentangle these explanations. Given the stronger associations with grey matter in ABCD and with white matter in IMAGEN, one possibility is that adversity influences distinct neurobiological mechanisms depending on the developmental window. The stronger associations with grey matter volume in ABCD may reflect earlier neuronal changes in grey matter (e.g., dendritic remodelling) between 9 and 12 years old (Bethlehem et al., 2022; Paus, 2023). In contrast, the stronger associations with white matter fractional anisotropy in IMAGEN could indicate later neuronal changes, such as myelination during mid-adolescence (12-15 years old) (Lebel et al., 2019; Paus, 2023).

Our analyses present limitations related to the choice of dimensionality reduction for the adversity components, the imaging methods, and the metrics used in this study. First, while probabilistic PCA worked well when questions were thematically coherent (e.g., Peer component captured 64% of the variance within the peer adversity items in IMAGEN), broader categories were less interpretable. For instance, Intrapersonal adversity included questions related to trauma experienced by the participant or in their family, from parental incarceration to accidents, yielding a less interpretable construct. This also resulted in limited comparability across cohorts, as Intrapersonal adversity items reflected different variables in each cohort, despite our efforts to construct comparable components. In contrast, results for the bullying and neighbourhood safety components were assessed using relatively similar items across cohorts, allowing for cautious comparison. Future research should incorporate more dimensions of adversity (e.g., societal-level adversity; see Rakesh et al., 2026), as well as use more standardised, harmonised adversity assessments, particularly in large-scale developmental studies.

Second, differences in scanner types, sites, and participant characteristics across and within the ABCD and IMAGEN cohorts may have introduced variability that reflects technical or sampling differences rather than true biological effects (Pan et al., 2024; Parsons et al., 2024). In addition, the distinction between grey and white matter metrics also depends on the metric itself, as measures capture overlapping neurobiological processes. For instance, cortical thickness has been shown to correlate with myelination in adjacent white matter (Natu et al., 2019). Future studies would benefit from harmonising imaging protocols to reduce methodological variability, as well as incorporating diverse structural metrics across both grey and white matter to better capture the range of biological processes influenced by adversity.

Our study assessed how adversity is differentially associated with grey and white matter trajectories using a longitudinal bivariate latent growth curve model in two large longitudinal cohorts. By showing that adversity is differentially associated with grey and white matter structures, and that these associations may vary by age, sex, and type of adversity, our findings emphasise the complexity of the relationship between adversity and brain development. This study contributes to a growing literature that underscores the importance of assessing multiple brain metrics in both grey and white matter across developmental stages and in relation to different forms of adversity. It also highlights the need to refine both our conceptual and measurement models of adversity to better capture its effects on the developing brain.

Beyond these theoretical contributions, our findings have practical implications. Identifying which types of adversity are associated with the greatest neural differences can help tailor intervention efforts to specific forms of adversity. In our study, neighbourhood safety showed the strongest associations with both grey matter volume (in ABCD) and fractional anisotropy (in IMAGEN). These neuronal differences possibly reflect cognitive adaptations developed in distinct neighbourhood contexts. Understanding these strategic adaptations enables the formulation of tailored interventions building more impactful policies (DeJoseph et al., 2024).

## Acknowledgements

R.A.K. and L.C.M. were supported by a Hypatia Fellowship at the RadboudUMC.

For the ABCD sample: Data used in the preparation of this article were obtained from the Adolescent Brain Cognitive Development™ (ABCD) Study, held in the NIH Brain Development Cohorts Data Sharing Platform. This is a multisite, longitudinal study designed to recruit more than 10,000 children aged 9–10 and follow them over 10 years into early adulthood. The ABCD Study® is supported by the National Institutes of Health and additional federal partners under award numbers: U01DA041048, U01DA050989, U01DA051016, U01DA041022, U01DA051018, U01DA051037, U01DA050987, U01DA041174, U01DA041106, U01DA041117, U01DA041028, U01DA041134, U01DA050988, U01DA051039, U01DA041156, U01DA041025, U01DA041120, U01DA051038, U01DA041148, U01DA041093, U01DA041089, U24DA041123, U24DA041147. A full list of supporters is available at Federal Partners – ABCD Study. ABCD Consortium investigators designed and implemented the study and/or provided data but did not necessarily participate in the analysis or writing of this report. This manuscript reflects the views of the authors and may not reflect the opinions or views of the NIH or ABCD Consortium investigators.

For the IMAGEN sample: This work received support from the following sources: the European Union-funded FP6 Integrated Project IMAGEN (Reinforcement-related behaviour in normal brain function and psychopathology) (LSHM-CT- 2007-037286), the Horizon 2020 funded ERC Advanced Grant ‘STRATIFY’ (Brain network based stratification of reinforcement-related disorders) (695313), Horizon Europe ‘environMENTAL’, grant no: 101057429, UK Research and Innovation (UKRI) Horizon Europe funding guarantee (10041392 and 10038599), Human Brain Project (HBP SGA 2, 785907, and HBP SGA 3, 945539), the Chinese government via the Ministry of Science and Technology (MOST). The German Center for Mental Health (DZPG), the Bundesministerium für Bildung und Forschung (BMBF grants 01GS08152; 01EV0711; Forschungsnetz AERIAL 01EE1406A, 01EE1406B; Forschungsnetz IMAC-Mind 01GL1745B), the Deutsche Forschungsgemeinschaft (DFG project numbers 458317126 [COPE], 186318919 [FOR 1617], 178833530 [SFB 940], 386691645 [NE 1383/14-1], 402170461 [TRR 265], 454245598 [IRTG 2773]), the Medical Research Foundation and Medical Research Council (grants MR/R00465X/1 and MR/S020306/1), the National Institutes of Health (NIH) funded ENIGMA-grants 5U54EB020403-05, 1R56AG058854-01 and U54 EB020403 as well as NIH R01DA049238, the National Institutes of Health, Science Foundation Ireland (16/ERCD/3797). NSFC grant 82150710554. Further support was provided by grants from: the Eranet Neuron (Grant ANR-18-NEUR00002-01– ADORe); Agence Nationale de la Recherche (Grant ANR-12-SAMA-0004 -GeBra); Assistance-Publique Hôpitaux-de-Paris and INSERM (interface grant); Paris Descartes University (Grant collaborative-project-2010); Paris Sud University (Grant IDEX-2012); Fondation de l’Avenir (Grant AP-RM-17-013); Fondation de France (Grant 00081242); Fédération pour la Recherche sur le Cerveau, and Fondation pour la Recherche Médicale (Grants DPA20140629802 and ADOLIMIS DPP20151033945); the Ile-de-France Region (Action 16700103 -grant to QIM– VEAVE, n°23002745–23002747).

