## Supplementary Material for "Adversity and adolescent brain development: differential associations with grey and white matter across two longitudinal cohorts"

### Contents


**I. Questionnaires used to form the four components of adversity**

1. ABCD
2. IMAGEN

**II. Component loadings for the four components of adversity**

**III. Results from latent class analyses and growth mixture models**

1. Cortical Grey Matter Volume
2. White Matter Fractional Anisotropy

**IV. Parameter estimates for the individual models**

1. ABCD
2. IMAGEN

**V. Leave one site out analyses**

1. ABCD
2. IMAGEN

**VI. Distribution of adversity between males and females**

1. ABCD
2. IMAGEN

**VII. Sex-stratified combined models for males and females**

1. ABCD
2. IMAGEN

### Questionnaires used to form the four components of adversity

#### ABCD

*Supplementary Table 1. Items and questionnaires details for each component of adversity used in the probabilistic PCA for ABCD*

| Adversity component | Items [name] | Questionnaire [name] Reported by |
|---|---|---|
| Intrapersonal | A car accident in which your child or another person in the car was hurt bad enough to require medical attention [ksads_ptsd_raw_754_p] | KSADS Diagnostic Interview for DSM-5 (Post-Traumatic Stress Disorder) - Parent [mh_p_ksads_ptsd] *Reported by the caregivers* *Measured at baseline* |
| | Another significant accident for which your child needed specialized and intensive medical treatment [ksads_ptsd_raw_755_p] | |
| | Witnessed or caught in a fire that caused significant property damage or personal injury [ksads_ptsd_raw_756_p] | |
| | Witnessed or caught in a natural disaster that caused significant property damage or personal injury [ksads_ptsd_raw_757_p] | |
| | Witnessed or present during an act of terrorism (e.g., Boston marathon bombing) [ksads_ptsd_raw_758_p] | |
| | Witnessed death or mass destruction in a war zone [ksads_ptsd_raw_759_p] | |
| | Witnessed someone shot or stabbed in the community [ksads_ptsd_raw_760_p] | |
| | Learned about the sudden unexpected death of a loved one [ksads_ptsd_raw_770_p] | |
| | At any time in your child's life, was anyone close to your child murdered, like a friend, neighbor or someone in your child's family? [socialdev_pvict_w6] | Victimization Questionnaire [sd_p_vict] *Reported by the caregivers* *Measured at baseline* |
| | At any time in your child's life, was your child in any place in real life where he/she could see or hear people being shot, bombs going off, or street riots? [socialdev_pvict_w8] | |
| | At any time in your child's life, was your child in the middle of a war where he/she could hear real fighting with guns or bombs? [socialdev_pvict_w9] | |
| | Someone in family died? [ple_died_y] | Negative Life Events Questionnaire* [mh_y_le] *Reported by the adolescent* *Measured at baseline* |
| | Family member was seriously injured? [ple_injured_y] | |
| | Saw crime or accident? [ple_crime_y] | |
| | Close friend was seriously sick/injured? [ple_friend_injur_y] | |
| | Family member had drug and/or alcohol problem? [ple_sud_y] | |
| | You got seriously sick? [ple_ill_y] | |
| | You got seriously injured? [ple_injur_y] | |
| | Mother/father figure lost job? [ple_job_y] | |
| | Someone in the family was arrested? [ple_arrest_y] | |
| | Close friend died? [ple_friend_died_y] | |
| | Brother or sister left home? [ple_sib_y] | |
| | Was a victim of crime/violence/assault? [ple_victim_y] | |
| | Parents separated or divorced? [ple_separ_y] | |
| | Parents/caregiver got into trouble with the law? [ple_law_y] | |
| | Attended a new school? [ple_school_y] | |
| | Family moved? [ple_move_y] | |
| | One of the parents/caregivers went to jail? [ple_jail_y] | |
| | Got new stepmother or stepfather? [ple_step_y] | |
| | Parent/caregiver got a new job? [ple_new_job_y] | |
| | Got new brother or sister? [ple_new_sib_y] | |
| Caregivers | Caregiver 1 makes me feel better after talking over my worries with him/her [crpbi_parent1_y] | |

| | | |
|---|---|---|
| | Caregiver 1 smiles at me very often [crpbi_parent2_y] | PhenX Neighborhood Safety/Crime Survey – Youth [ce_y_crpbi] *Reported by the adolescent Measured at baseline* |
| | Caregiver 1 is able to make me feel better when I am upset [crpbi_parent3_y] | |
| | Caregiver 1 believes in showing his/her love for me [crpbi_parent4_y] | |
| | Caregiver 1 is easy to talk to [crpbi_parent5_y] | |
| | Caregiver 2 makes me feel better after talking over my worries with him/her [crpbi_caregiver12_y] | |
| | Caregiver 2 smiles at me very often [crpbi_caregiver13_y] | |
| | Caregiver 2 is able to make me feel better when I am upset [crpbi_caregiver14_y] | |
| | Caregiver 2 believes in showing his/her love for me [crpbi_caregiver15_y] | |
| | Caregiver 2 is easy to talk to [crpbi_caregiver16_y] | |
| **Peers** | Do you have any problems with bullying at school or in your neighborhood? [ksads_bully_raw_26] | KSADS Background Items Survey – Youth [mh_y_ksads_bg] *Reported by the caregivers Measured at baseline* |
| | Sometimes groups of kids or gangs attack people. At any time in your life, did a group of kids or a gang hit, jump, or attack you? [socialdev_cvict_p1] | Victimization Questionnaire [sd_y_vict] *Reported by the adolescents Measured at baseline* |
| | At any time in your life, did any kid, even a brother or sister, hit you? Somewhere like at home, at school, out playing, in a store, or anywhere else? [socialdev_cvict_p2] | |
| | At any time in your life, did any kids try to hurt your private parts on purpose by hitting or kicking you there? [socialdev_cvict_p3] | |
| | At any time in your life, did any kids, even a brother or sister, pick on you by chasing or grabbing you, or by making you do something you didn't want to do? [socialdev_cvict_p4] | |
| | At any time in your life, did you get really scared or feel really bad because kids were calling you names, saying mean things to you, or saying they didn't want you around? [socialdev_cvict_p5] | |
| | At any time in your life, did a boyfriend or girlfriend or anyone you went on a date with slap or hit you? [socialdev_cvict_p6] | |
| | At any time in your life, did any kids ever tell lies or spread rumors about you or try to make others dislike you? [socialdev_cvict_p7] | |
| | At any time in your life, did any kids ever keep you out of things on purpose, exclude you from their group of friends, or completely ignore you? [socialdev_cvict_p8] | |
| | Has anyone ever used the internet to bother or harass you or to spread mean words or pictures about you? [socialdev_cvict_int1] | |
| | Has anyone ever used a cell phone or texting to bother or harass you or to spread mean words or pictures about you? [socialdev_cvict_int2] | |
| | Lost a close friend? [ple_friend_y] | Negative Life Events Questionnaire* [mh_y_le] *Reported by the adolescent Measured at baseline* |
| **Community** | I feel safe walking in my neighborhood, day or night. [neighborhood1r_p] | PhenX Neighborhood Safety/Crime Survey - Parent [ce_p_nsc] *Reported by the caregivers* |
| | Violence is not a problem in my neighborhood. [neighborhood2r_p] | |
| | My neighborhood is safe from crime. [neighborhood3r_p] | |

| | | |
|---|---|---|
| | | *Measured at baseline* |
| | My neighborhood is safe from crime. [neighborhood_crime_y] | PhenX Neighborhood Safety/Crime Survey - Youth [ce_y_nsc] *Reported by the adolescent Measured at baseline* |

*Notes: For the items from the Negative Life Events Questionnaire we also included the questions assessing whether this was a good or bad experience (1=Mostly good;2=Mostly bad;6=Not applicable;7=Don't know) and how much did the event affect them (0=Not at All;1=A Little;2=Some;3=A lot). If the answers for those questions were 0 or 1, we replaced the items responses by 0.*

### IMAGEN

*Supplementary Table 2. Items and questionnaires details for each component of adversity used in the probabilistic PCA for IMAGEN*

| **Adversity component** | **Items [name]** | **Questionnaire [name] Reported by** |
|---|---|---|
| **Intrapersonal** | Parents divorced [leq_01_ever] | Life-Events Questionnaire [leq_T1] *Reported by the adolescent Measured at baseline* |
| | Family accident or illness [leq_02_ever] | |
| | Got in trouble with the law [leq_04_ever] | |
| | Death in family [leq_08_ever] | |
| | Serious accident or illness [leq_37_ever] | |
| | PTSD: Serious accident [p1e2a] | The Development and Well-Being Assessment Interview [dawba] *Reported by the caregivers Measured at baseline* |
| | PTSD: Witnessed attack [p1e2i] | |
| | PTSD: Witnessed accident, sudden death [p1e2j] | |
| | Serious accident [p1recentle1] | |
| | In hospital for serious illness [p1recentle2] | |
| | Death of parent, sibling or friend [p1recentle3] | |
| | Parental separation [p1recentle6] | |
| | Other stressful life event [p1recentle7] | |
| **Caregivers** | My mother was not very affectionate. [PAAQ_3] | Childhood Relationships Questionnaire [ce_y_crpbi] *Reported by the adolescent Measured at the second timepoints (19 years old)* |
| | When I was a young child and little things went wrong I did not feel sure I could count on my mother to take care of me. [PAAQ_4] | |
| | When I was a child my mother sometimes told me that if I was not good she would stop loving me. [PAAQ_10] | |
| | In childhood I knew I was low on my mother's priority list. [PAAQ_18] | |
| | ] In childhood my mother sometimes threatened to leave me or to send me away if I wasn't good. [PAAQ_23] | |
| | In childhood I often had the impression that my mother was not listening to me. She often tuned me out. [PAAQ_29] | |
| | If something really bad happened to me in childhood I did not feel I could count on my mother to support me. [PAAQ_45] | |
| | When I was a child I sometimes got the feeling that my mother wished I was never born. [PAAQ_46] | |
| | In childhood my mother often told me she was sacrificing herself for me. [PAAQ_49] | |
| | When I acted bad as a child my mother would, at times, threaten to send me away. [PAAQ_52] | |
| | I never felt like my mother gave me enough attention. [PAAQ_53] | |
| | Non-physical punishment [p1flq16] | |
| | Gets love and affection [p1flq10] | |

| | | |
|---|---|---|
| | Gets blamed unfairly [p1flq19] | The Development and Well-Being Assessment Interview [dawba] *Reported by the caregivers* *Measured at baseline* |
| **Peers** | I was bullied at school (a student/ peer said or did nasty or unpleasant things to me).. [C.bully01] | Bully Questionnaire [bully] *Reported by the adolescent* *Measured at baseline* |
| | I was called mean names, was made fun of, or teased in a hurtful way by a student/ peer. [C.bully02] | |
| | A student/ peer left me out of things on purpose, excluded me from their group of friends or completely ignored me. [C.bully03] | |
| | I was hit, kicked, pushed or shoved around, or locked indoors by a student/ peer. [C.bully04] | |
| | Loss of close friendship [p1recentle4] | The Development and Well-Being Assessment Interview [dawba] *Reported by the caregivers* *Measured at baseline* |
| **Community** | Family stresses: Neighbours or neighbourhood [p1fs5] | The Development and Well-Being Assessment Interview [dawba] *Reported by the caregivers* *Measured at baseline* |

### Component loadings for the four components of adversity

*Supplementary Table 3. Items for each component of adversity used in the probabilistic PCA and the amount of variance explained by items for ABCD and IMAGEN. [C] is an item responded by the primary caregiver and [A] is an item responded by the adolescent.*

| | ABCD | | IMAGEN | |
|---|---|---|---|---|
| | *Items* | *Component loadings* | *Items* | *Component loadings* |
| **Intrapersonal adversity** | [C] Car accident requiring medical attention | 0.10 | Parents divorced [A] | 0.45 |
| | [C] Other significant accident requiring medical treatment for child | 0.06 | Family accident or illness [A] | 0.42 |
| | [C] Witnessed or caught in a fire that caused significant property damage or personal injury | 0.07 | Got in trouble with the law [A] | 0.18 |
| | [C] Witnessed or caught in a natural disaster that caused significant property damage or personal injury | 0.06 | Death in family [A] | 0.25 |
| | [C] Witnessed or present during an act of terrorism | 0.07 | Serious accident or illness [A] | 0.33 |
| | [C] Witnessed death or mass destruction in a war zone | 0.09 | [C] Serious accident (PTSD) | 0.001 |
| | [C] Witnessed someone shot or stabbed in the community | 0.12 | [C] Witnessed attack (PTSD) | 0.01 |
| | [C] Sudden unexpected death of a loved one | 0.13 | [C] Witnessed accident, sudden death (PTSD) | 0.02 |
| | [C] Anyone close to your child murdered, like a friend, neighbor or someone in your child's family | 0.04 | [C] Serious accident | 0.12 |
| | [C] Child in any place in real life where they could see or hear people being shot, bombs going off, or street riots | 0.05 | [C] In hospital for serious illness | 0.16 |
| | [C] Child in the middle of a war where they could hear real fighting with guns or bombs | 0.03 | [C] Death of parent, sibling or friend | 0.20 |
| | Someone in family died [A] | 0.18 | [C] Parental separation | 0.46 |
| | Family member seriously injured [A] | 0.24 | [C] Other stressful life event | 0.34 |
| | Saw crime or accident [A] | 0.18 | | |
| | Close friend seriously sick/injured [A] | 0.17 | | |
| | Family member had drug and/or alcohol problem [A] | 0.26 | | |
| | Got seriously sick [A] | 0.21 | | |
| | Got seriously injured [A] | 0.19 | | |
| | Mother/father figure lost job [A] | 0.18 | | |
| | Someone in the family arrested [A] | 0.39 | | |
| | Close friend died [A] | 0.09 | | |
| | Brother or sister left home [A] | 0.15 | | |
| | Victim of crime/violence/assault [A] | 0.10 | | |
| | Parents separated or divorced [A] | 0.23 | | |

| | | | | |
|---|---|---|---|---|
| | Trouble with the law for caregiver [A] | 0.34 | | |
| | Attended a new school [A] | 0.14 | | |
| | Family moved [A] | 0.15 | | |
| | One of the caregivers went to jail [A] | 0.39 | | |
| | Got new stepmother or stepfather [A] | 0.18 | | |
| | Caregiver got a new job [A] | 0.07 | | |
| | Got new brother or sister [A] | 0.04 | | |
| **Caregiver adversity** | Caregiver 1 makes me feel better after talking over my worries with them [inverted] [A] | 0.31 | My mother was not very affectionate [A] | 0.30 |
| | Caregiver 1 smiles at me very often [inverted] [A] | 0.30 | When I was a young child and little things went wrong I did not feel sure I could count on my mother to take care of me. [A] | 0.25 |
| | Caregiver 1 is able to make me feel better when I am upset [inverted] [A] | 0.33 | When I was a child my mother sometimes told me that if I was not good she would stop loving me. [A] | 0.30 |
| | Caregiver 1 believes in showing their love for me [inverted] [A] | 0.28 | In childhood I knew I was low on my mother's priority list. [A] | 0.27 |
| | Caregiver 1 is easy to talk to [inverted] [A] | 0.29 | In childhood my mother sometimes threatened to leave me or to send me away if I wasn't good. [A] | 0.32 |
| | Caregiver 2 makes me feel better after talking over my worries with them [inverted] [A] | 0.35 | In childhood I often had the impression that my mother was not listening to me. She often tuned me out. [A] | 0.33 |
| | Caregiver 2 smiles at me very often [inverted] [A] | 0.32 | If something really bad happened to me in childhood I did not feel I could count on my mother to support me. [A] | 0.25 |
| | Caregiver 2 is able to make me feel better when I am upset [inverted] [A] | 0.36 | When I was a child I sometimes got the feeling that my mother wished I was never born. [A] | 0.34 |
| | Caregiver 2 believes in showing their love for me [inverted] [A] | 0.32 | In childhood my mother often told me she was sacrificing herself for me. [A] | 0.20 |
| | Caregiver 2 is easy to talk to [inverted] [A] | 0.30 | When I acted bad as a child my mother would, at times, threaten to send me away. [A] | 0.32 |
| | | | I never felt like my mother gave me enough attention. [A] | 0.28 |
| | | | [C] Non-physical punishment | 0.15 |
| | | | [C] Gets love and affection [inverted] | 0.20 |
| | | | [C] Gets blamed unfairly | 0.08 |
| **Peer** | Problems with bullying at school or in your neighborhood [A] | 0.79 | Bullied at school (a student/ peer said or did nasty or | 0.56 |

| | | | | |
|---|---|---|---|---|
| **adversity** | | | unpleasant things to me). [A] | |
| | Group of kids or a gang hit, jump, or attack you [A] | 0.06 | Called mean names, was made fun of, or teased in a hurtful way by a student/ peer. [A] | 0.54 |
| | Kid, brother or sister hit you [A] | 0.03 | A student/ peer left me out of things on purpose, excluded me from their group of friends or completely ignored me. [A] | 0.44 |
| | Kids try to hurt your private parts on purpose by hitting or kicking you there [A] | 0.05 | Hit, kicked, pushed or shoved around, or locked indoors by a student/ peer. [A] | 0.43 |
| | Kids, even a brother or sister, pick on you by chasing or grabbing you, or by making you do something you didn't want to do [A] | 0.03 | [C] Loss of close friendship | 0.14 |
| | Really scared or feel really bad because kids were calling you names, saying mean things to you, or saying they didn't want you around [A] | 0.09 | | |
| | Boyfriend or girlfriend or anyone you went on a date with slap or hit you [A] | 0.02 | | |
| | Kids ever tell lies or spread rumors about you or try to make others dislike you [A] | 0.08 | | |
| | Kids ever keep you out of things on purpose, exclude you from their group of friends, or completely ignore you [A] | 0.08 | | |
| | Kids bother or harass or spread mean words/pictures about you on the internet [A] | 0.06 | | |
| | Has anyone ever used a cell phone or texting to bother or harass you or to spread mean words or pictures about you [A] | 0.06 | | |
| | Lost a close friend [A] | 0.58 | | |
| **Community adversity** | [C] I feel safe walking in my neighborhood, day or night [inverted] | 0.54 | [C] Family stresses: Neighbours or neighbourhood | |
| | [C] Violence is not a problem in my neighborhood [inverted] | 0.57 | | |
| | [C] My neighborhood is safe from crime [inverted] | 0.56 | | |
| | My neighborhood is safe from crime [inverted] [A] | 0.28 | | |

### Results from latent class analyses and growth mixture models

#### Cortical Grey Matter Volume

*Supplementary Table 4. Comparison of the Latent Class Growth Analyses for 1, 2, 3 and 4 groups for total cortical grey matter volume in ABCD*

| Number of groups | Log likelihood | Number of parameters | BIC | % class 1 | % class 2 | % class 3 | % class 4 |
|---|---|---|---|---|---|---|---|
| 1 | -282663.7 | 3 | 565355.6 | 100 | | | |
| 2 | -278292.9 | 6 | 556642.0 | 43.7 | 56.3 | | |
| 3 | -275445.4 | 9 | 550975.1 | 20.1 | 29.4 | 50.5 | |
| 4 | -282536.9 | 12 | 565186.3 | 0 | 50.9 | 0 | 49.1 |

*Supplementary Figure 1. Trajectories of total cortical grey matter for ABCD. The colors represent the three groups computed through the Latent Class Growth Analysis.*

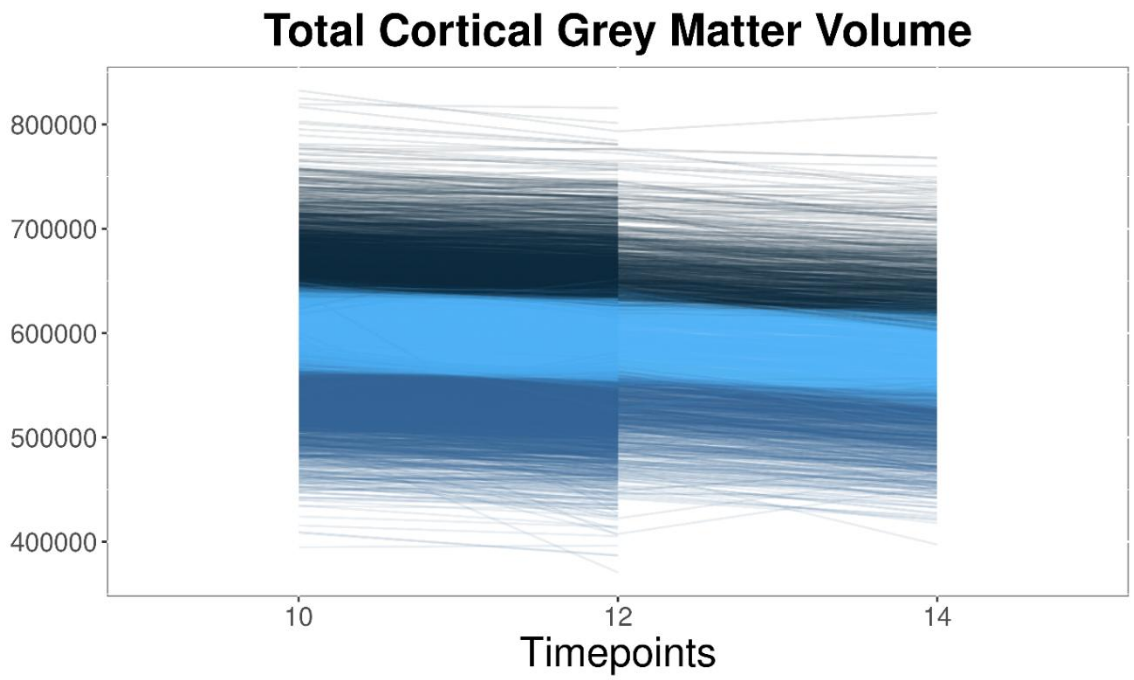


*Supplementary Table 5. Comparison of the Latent Class Analyses for 1, 2, 3 and 4 groups for total cortical grey matter volume in IMAGEN*

| Number of groups | Log likelihood | Number of parameters | BIC | % class 1 | % class 2 | % class 3 | % class 4 |
|---|---|---|---|---|---|---|---|
| 1 | -36634.6 | 3 | 73289.8 | 100 | | | |
| 2 | -3.6136.4 | 6 | 72314.1 | 55.3 | 44.7 | | |
| 3 | -3.5885.5 | 9 | 71833.1 | 14.5 | 32.9 | 52.6 | |
| 4 | -1000000000 | 12 | 2000000000 | 0 | 0 | 0 | 0 |

*Supplementary Figure 2. Trajectories of total cortical grey matter for IMAGEN. The colors represent the three groups computed through the Latent Class Growth Analysis.*

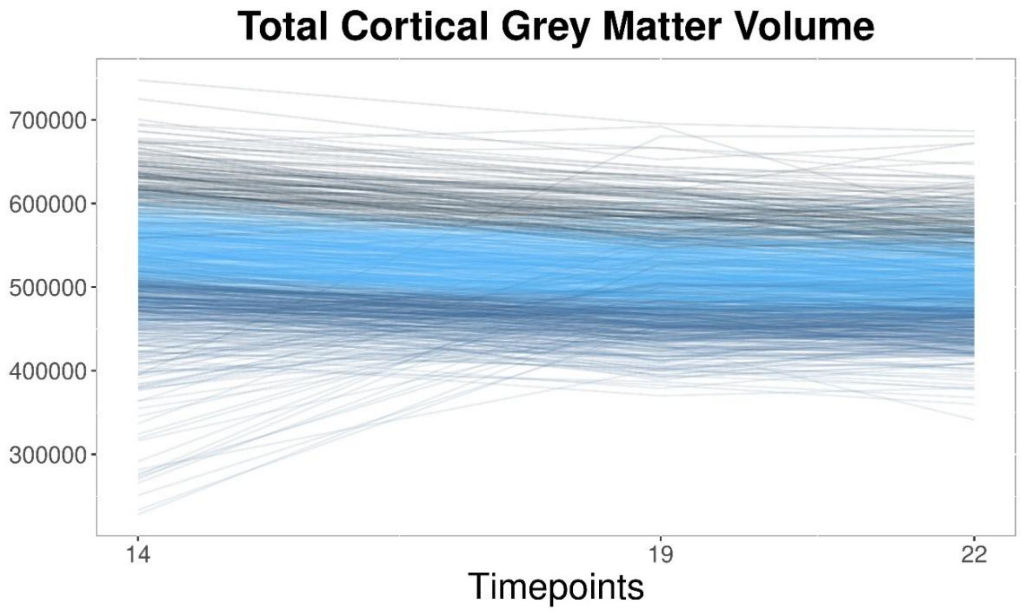


### White Matter Fractional Anisotropy

*Supplementary Table 6. Comparison of the Latent Class Analyses for 1, 2, 3 and 4 groups for mean fractional anisotropy in ABCD*

| Number of groups | Log likelihood | Number of parameters | BIC | % class 1 | % class 2 | % class 3 | % class 4 |
|---|---|---|---|---|---|---|---|
| 1 | 51958.40 | 3 | -103888.7 | 100 | | | |
| 2 | 54403.89 | 6 | -108751.7 | 17.8 | 82.2 | | |
| 3 | 55088.20 | 9 | -110092.2 | 11.1 | 43.9 | 45 | |
| 4 | 55425.26 | 12 | -110738.3 | 14.5 | 55 | 2 | 28.5 |

*Supplementary Figure 3. Trajectories of mean fractional anisotropy for ABCD. The colors represent the three groups computed through the Latent Class Growth Analysis.*

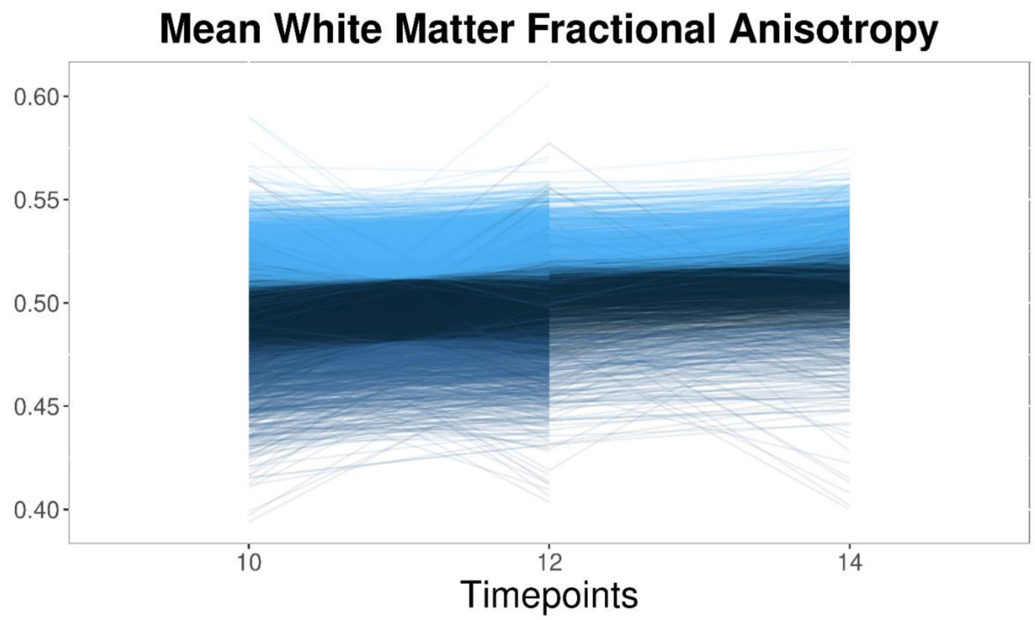


*Supplementary Table 7. Comparison of the Latent Class Analyses for 1, 2, 3 and 4 groups for mean fractional anisotropy in IMAGEN*

| Number of groups | Log likelihood | Number of parameters | BIC | % class 1 | % class 2 | % class 3 | % class 4 |
|---|---|---|---|---|---|---|---|
| 1 | 8361.9 | 3 | -16701.6 | 100 | | | |
| 2 | 8898.3 | 6 | -17751.8 | 44.9 | 55.1 | | |
| 3 | 9174.3 | 9 | -18281.5 | 23.4 | 26 | 50.6 | |
| 4 | 9298.4 | 12 | -18507.3 | 8.9 | 37.5 | 36.7 | 16.9 |

*Supplementary Figure 4. Trajectories of mean fractional anisotropy for IMAGEN. The colors represent the three groups computed through the Latent Class Growth Analysis.*

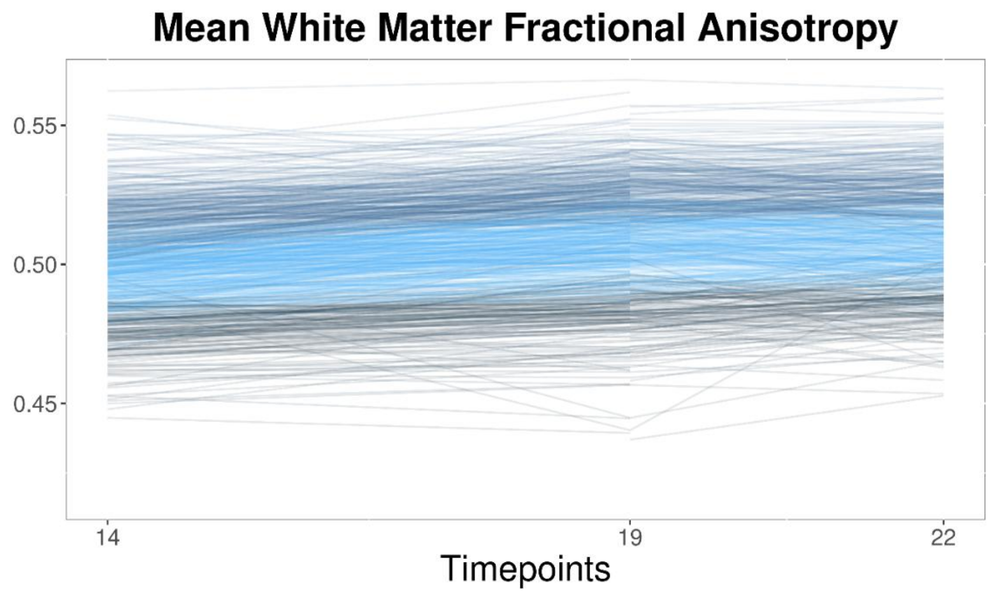

### Parameter estimates for the individual models

#### ABCD

*Supplementary Table 8. Parameters estimates of the bivariate latent growth curve model with only Intrapersonal adversity predicting the intercepts and slopes of total cortical grey matter volume and mean fractional anisotropy in ABCD.*

**Individual model with Intrapersonal adversity (ABCD)**

*Fit indices: $x^2$ = 158.096, df= 12, p<0.001, RMSEA = 0.060, CFI= 0.996, TLI=0.992, SRMR= 0.014*

| Parameters | Estimate | Std.error | z-value | p-value | Std.estimate | CI lower | CI upper |
|---|---|---|---|---|---|---|---|
| **Intrapersonal** | | | | | | | |
| Intercept GMV | -638 | 496 | -1.29 | 0.199 | -0.013 | -1610 | 334 |
| Intercept FA | -7.55E-05 | 1.92E-04 | -0.39 | 0.693 | -0.005 | -4.51E-04 | 3.00E-04 |
| Slope GMV | 174 | 229 | 0.76 | 0.448 | 0.018 | -275 | 622 |
| Slope FA | -1.97E-05 | 7.55E-05 | -0.26 | 0.794 | -0.007 | -1.68E-04 | 1.28E-04 |

*Supplementary Table 9. Parameters estimates of the bivariate latent growth curve model with only Caregivers adversity predicting the intercepts and slopes of total cortical grey matter volume and mean fractional anisotropy in ABCD.*

**Individual model with Caregiver adversity (ABCD)**

*Fit indices: $x^2$ = 147.539, df= 12, p<0.001, RMSEA = 0.058, CFI= 0.996, TLI=0.993, SRMR= 0.014*

| Parameters | Estimate | Std.error | z-value | p-value | Std.estimate | CI lower | CI upper |
|---|---|---|---|---|---|---|---|
| **Caregiver** | | | | | | | |
| Intercept GMV | -884 | 515 | -1.72 | 0.086 | -0.019 | -1894 | 126 |
| Intercept FA | -5.25E-04 | 1.84E-04 | -2.86 | ***0.004*** | -0.032 | -8.85E-04 | -1.66E-04 |
| Slope GMV | 354 | 231 | 1.53 | 0.125 | 0.039 | -98 | 807 |
| Slope FA | 4.08E-05 | 8.81E-05 | 0.46 | 0.643 | 0.016 | -1.32E-04 | 2.14E-04 |

*Supplementary Table 10. Parameters estimates of the bivariate latent growth curve model with only Peers adversity predicting the intercepts and slopes of total cortical grey matter volume and mean fractional anisotropy in ABCD.*

**Individual model with Peer adversity (ABCD)**

| *Fit indices: x² = 147.351, df= 12, p<0.001, RMSEA = 0.058, CFI= 0.996, TLI=0.993, SRMR= 0.014* | | | | | | | |
|---|---|---|---|---|---|---|---|
| Parameters | Estimate | Std.error | z-value | p-value | Std.estimate | CI lower | CI upper |
| **Peer** | | | | | | | |
| Intercept GMV | -4164 | 1040 | -4.01 | ***<0.001*** | -0.042 | -6202 | -2126 |
| Intercept FA | -2.26E-04 | 3.83E-04 | -0.59 | 0.555 | -0.007 | -9.76E-04 | 5.24E-04 |
| Slope GMV | 43 | 510 | 0.08 | 0.933 | 0.002 | -957 | 1043 |
| Slope FA | 1.10E-04 | 1.87E-04 | 0.59 | 0.557 | 0.020 | -2.57E-04 | 4.77E-04 |

*Supplementary Table 11. Parameters estimates of the bivariate latent growth curve model with only Community adversity predicting the intercepts and slopes of total cortical grey matter volume and mean fractional anisotropy in ABCD.*

**Individual model with Community adversity (ABCD)**

| *Fit indices: x² = 153.867, df= 12, p<0.001, RMSEA = 0.059, CFI= 0.996, TLI=0.992, SRMR= 0.014* | | | | | | | |
|---|---|---|---|---|---|---|---|
| Parameters | Estimate | Std.error | z-value | p-value | Std.estimate | CI lower | CI upper |
| **Community** | | | | | | | |
| Intercept GMV | -6165 | 581 | -10.61 | ***<0.001*** | -0.108 | -7303 | -5026 |
| Intercept FA | -1.34E-03 | 2.14E-04 | -6.27 | ***<0.001*** | -0.067 | -1.76E-03 | -9.22E-04 |
| Slope GMV | -60 | 289 | -0.21 | 0.836 | -0.005 | -626 | 506 |
| Slope FA | 1.85E-04 | 1.06E-04 | 1.75 | 0.080 | 0.059 | -2.22E-05 | 3.92E-04 |

*Supplementary Figure 5. Regression between the components of adversity and grey matter volume of fractional anisotropy for the significant relationships in ABCD.*

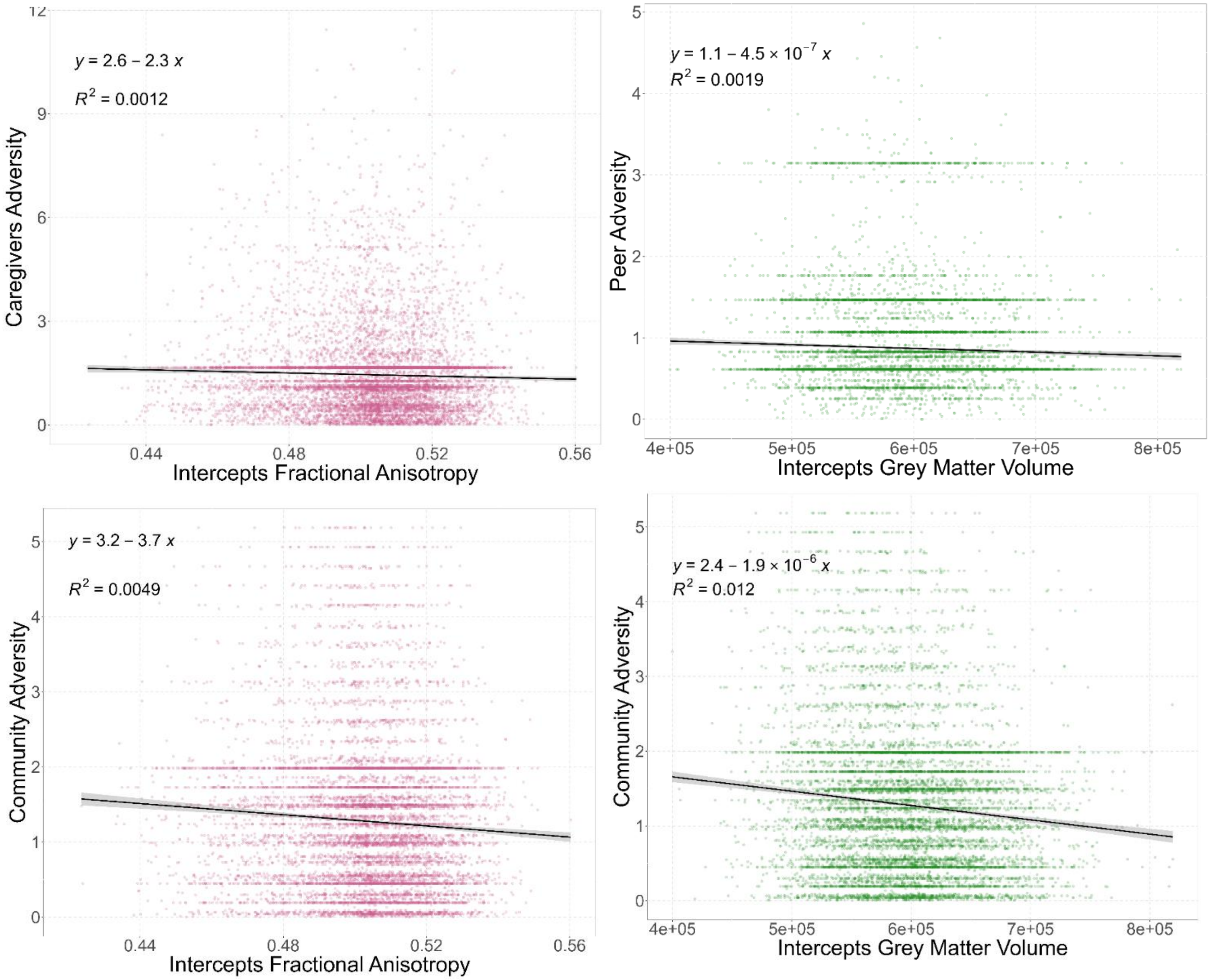

### IMAGEN

*Supplementary Table 12. Parameters estimates of the bivariate latent growth curve model with only Intrapersonal adversity predicting the intercepts and slopes of total cortical grey matter volume and mean fractional anisotropy in IMAGEN.*

**Individual model with Intrapersonal adversity (IMAGEN)**

*Fit indices: $x^2$ = 68.591, df= 11, p<0.001, RMSEA = 0.083, CFI= 0.987, TLI=0.975, SRMR= 0.022*

| Parameters | Estimate | Std.error | z-value | p-value | Std.estimate | CI lower | CI upper |
|---|---|---|---|---|---|---|---|
| **Intrapersonal** | | | | | | | |
| Intercept GMV | -2164 | 1945 | -1.11 | 0.266 | -0.041 | -5976 | 1648 |
| Intercept FA | 3.01E-04 | 3.92E-04 | 0.77 | 0.443 | 0.020 | -4.67E-04 | 1.07E-03 |
| Slope GMV | 2933 | 1631 | 1.80 | 0.072 | 0.080 | -265 | 6131 |
| Slope FA | 2.83E-04 | 2.97E-04 | 0.95 | 0.340 | 0.060 | -2.99E-04 | 8.64E-04 |

*Supplementary Table 13. Parameters estimates of the bivariate latent growth curve model with only Caregivers adversity predicting the intercepts and slopes of total cortical grey matter volume and mean fractional anisotropy in IMAGEN.*

**Individual model with Caregiver adversity (IMAGEN)**

*Fit indices: $x^2$ = 72.641, df= 11, p<0.001, RMSEA = 0.083, CFI= 0.987, TLI=0.974, SRMR= 0.022*

| Parameters | Estimate | Std.error | z-value | p-value | Std.estimate | CI lower | CI upper |
|---|---|---|---|---|---|---|---|
| **Caregiver** | | | | | | | |
| Intercept GMV | -760 | 925 | -0.82 | 0.411 | -0.023 | -2573 | 1053 |
| Intercept FA | -1.21E-03 | 2.78E-04 | -4.37 | ***<0.001*** | -0.130 | -1.76E-03 | -6.69E-04 |
| Slope GMV | 837 | 607 | 1.38 | 0.168 | 0.037 | -352 | 2027 |
| Slope FA | 1.56E-04 | 1.63E-04 | 0.96 | 0.339 | 0.053 | -1.63E-04 | 4.75E-04 |

*Supplementary Table 14. Parameters estimates of the bivariate latent growth curve model with only Peers adversity predicting the intercepts and slopes of total cortical grey matter volume and mean fractional anisotropy in IMAGEN.*

**Individual model with Peer adversity (IMAGEN)**

*Fit indices: $x^2$ = 73.761, df= 11, p<0.001, RMSEA = 0.085, CFI= 0.986, TLI=0.974, SRMR= 0.022*

| Parameters | Estimate | Std.error | z-value | p-value | Std.estimate | CI lower | CI upper |
|---|---|---|---|---|---|---|---|
| **Peer** | | | | | | | |
| Intercept GMV | -366 | 1402 | -0.26 | 0.794 | -0.009 | -3114 | 2383 |

| | | | | | | | |
|---|---|---|---|---|---|---|---|
| Intercept FA | -1.25E-03 | 3.04E-04 | -4.10 | ***<0.001*** | -0.103 | -1.84E-03 | -6.51E-04 |
| Slope GMV | 1061 | 961 | 1.10 | 0.270 | 0.036 | -823 | 2945 |
| Slope FA | 4.57E-04 | 2.63E-04 | 1.74 | 0.082 | 0.119 | -5.80E-05 | 9.72E-04 |

*Supplementary Table 15. Parameters estimates of the bivariate latent growth curve model with only Community adversity predicting the intercepts and slopes of total cortical grey matter volume and mean fractional anisotropy in IMAGEN.*

**Individual model with Community adversity (IMAGEN)**

*Fit indices: $x^2$ = 74.008, df= 11, p<0.001, RMSEA = 0.084, CFI= 0.986, TLI=0.974, SRMR= 0.022*

| Parameters | Estimate | Std.error | z-value | p-value | Std.estimate | CI lower | CI upper |
|---|---|---|---|---|---|---|---|
| **Community** | | | | | | | |
| Intercept GMV | -366 | 1402 | -0.26 | 0.794 | -0.009 | -3114 | 2383 |
| Intercept FA | -1.25E-03 | 3.04E-04 | -4.10 | ***<0.001*** | -0.103 | -1.84E-03 | -6.51E-04 |
| Slope GMV | 1061 | 961 | 1.10 | 0.270 | 0.036 | -823 | 2945 |
| Slope FA | 4.57E-04 | 2.63E-04 | 1.74 | 0.082 | 0.119 | -5.80E-05 | 9.72E-04 |

*Supplementary Figure 6. Regression between the components of adversity and grey matter volume of fractional anisotropy for the significant relationships in IMAGEN*

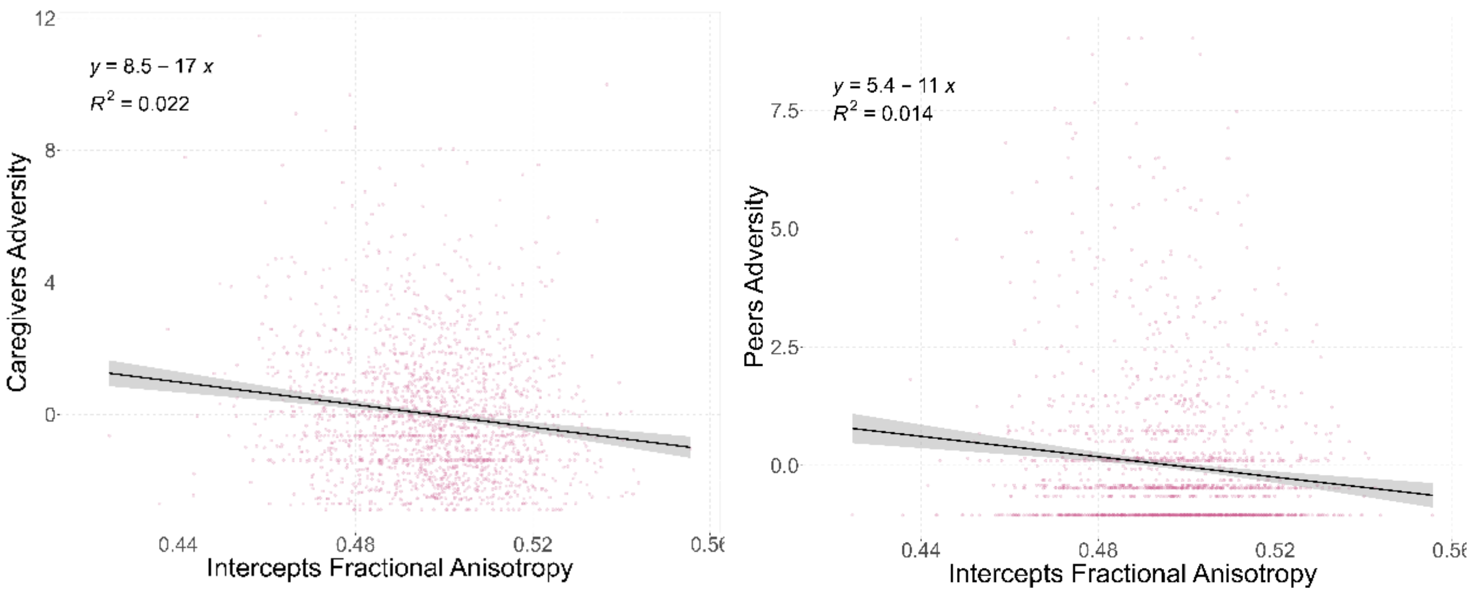

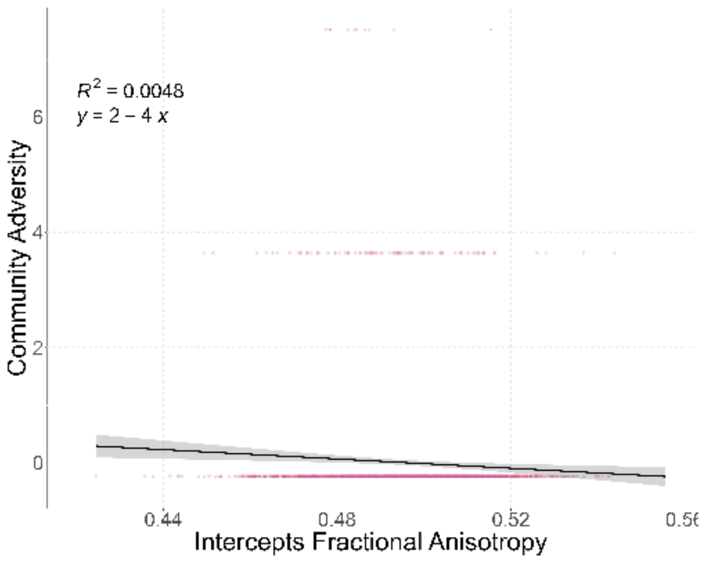


### Leave one site out analyses

#### ABCD

*Supplementary Figure 7. Leave one site out analyses in the full model with the four adversity in ABCD. Each dot represents the standard estimates found between one type of adversity and one neuroimaging variable. The color shows if the association was significant (red) or not. The dashed line shows the standard estimate found in the full model with every sites.*

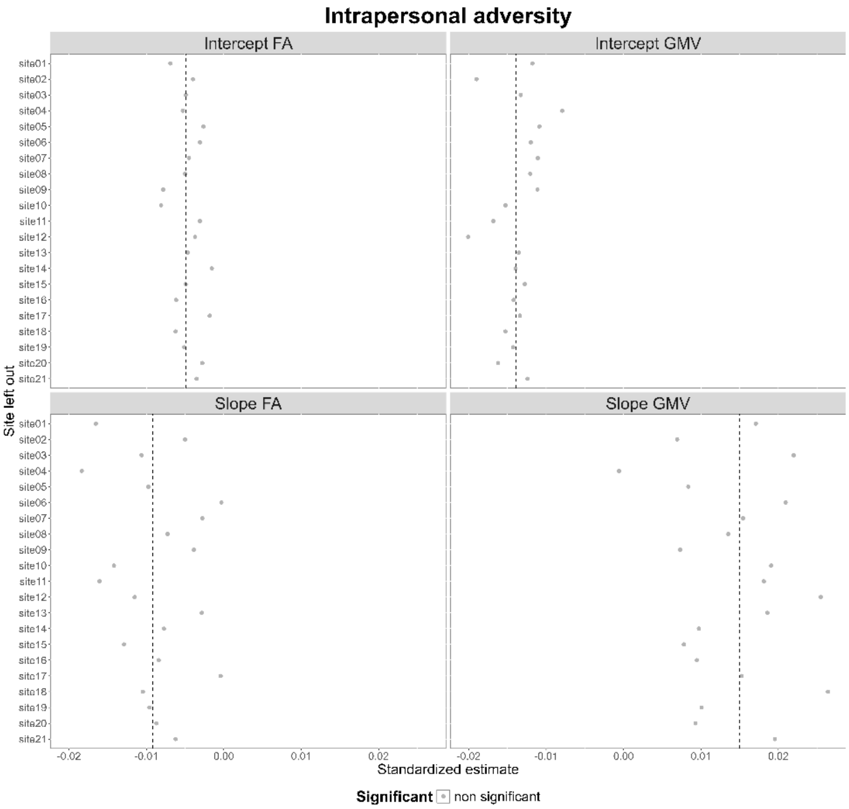
Intrapersonal adversity
Intercept FA
Intercept GMV
Slope FA
Slope GMV
Site left out
site01
site02
site03
site04
site05
site06
site07
site08
site09
site10
site11
site12
site13
site14
site15
site16
site17
site18
site19
site20
site21
-0.02
-0.01
0.00
0.01
0.02
Standardized estimate
Significant
non significant

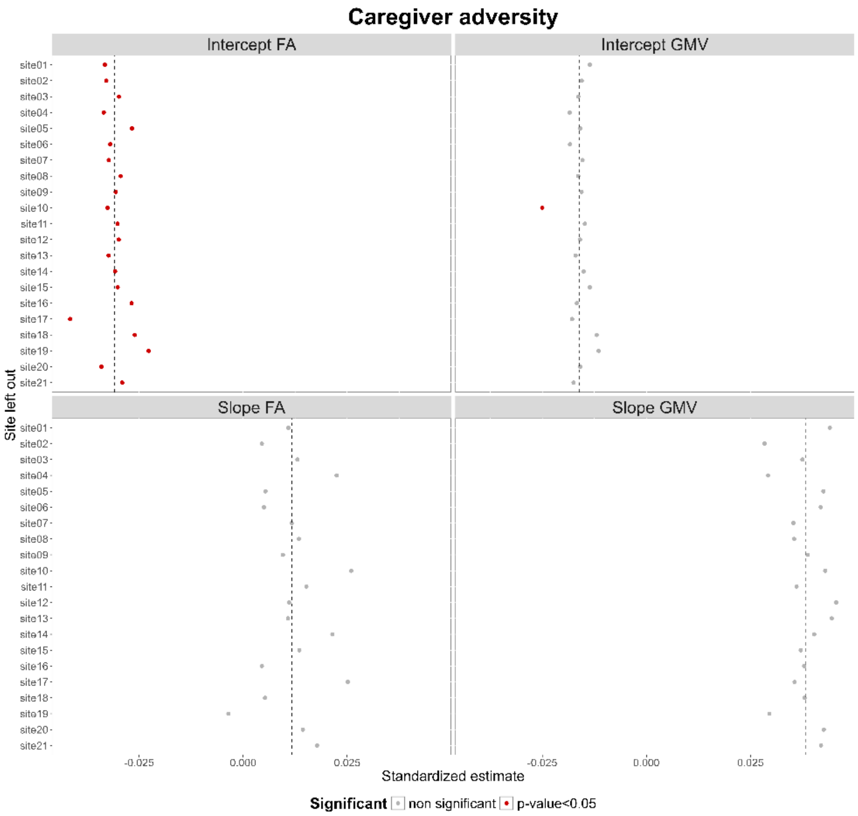
Caregiver adversity
Intercept FA
Intercept GMV
Slope FA
Slope GMV
Site left out
site01
site02
site03
site04
site05
site06
site07
site08
site09
site10
site11
site12
site13
site14
site15
site16
site17
site18
site19
site20
site21
-0.025
0.000
0.025
Standardized estimate
Significant
non significant
p-value<0.05

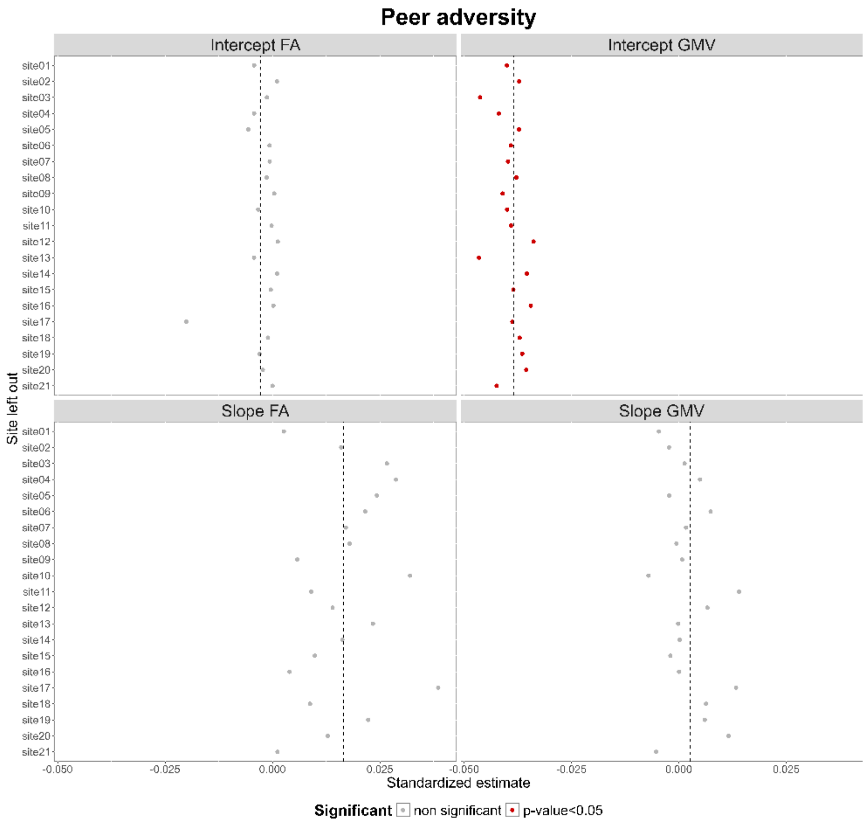
Peer adversity
Intercept FA
Intercept GMV
Slope FA
Slope GMV
Site left out
site01
site02
site03
site04
site05
site06
site07
site08
site09
site10
site11
site12
site13
site14
site15
site16
site17
site18
site19
site20
site21
-0.050
-0.025
0.000
0.025
Standardized estimate
Significant
non significant
p-value<0.05

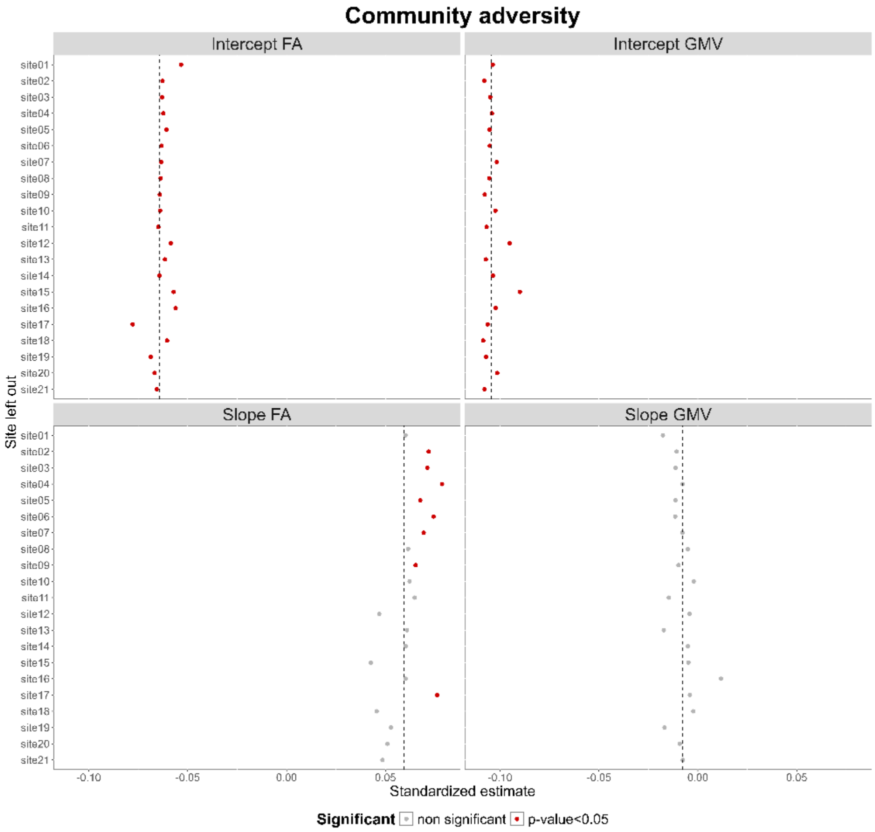
Community adversity
Intercept FA
Intercept GMV
Slope FA
Slope GMV
Site left out
site01
site02
site03
site04
site05
site06
site07
site08
site09
site10
site11
site12
site13
site14
site15
site16
site17
site18
site19
site20
site21
-0.10
-0.05
0.00
0.05
Standardized estimate
Significant
non significant
p-value<0.05

### IMAGEN

*Supplementary Figure 8. Leave one site out analyses in the full model with the four adversity in IMAGEN. Each dot represents the standard estimates found between one type of adversity and one neuroimaging variable. The color shows if the association was significant (red) or not. The dashed line shows the standard estimate found in the full model with every sites.*

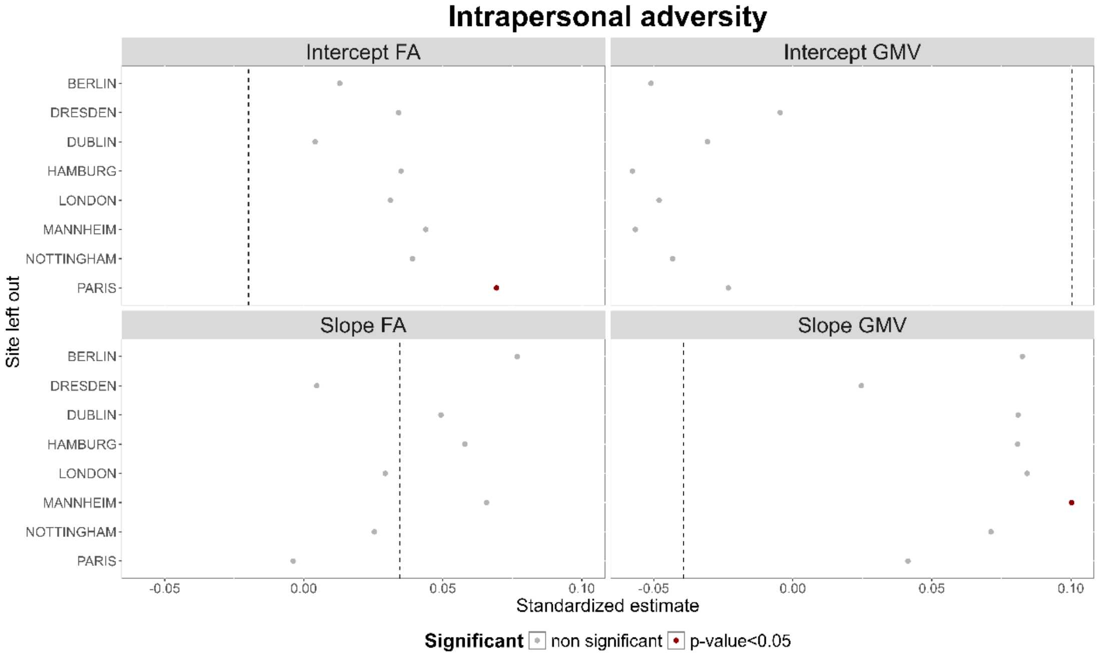


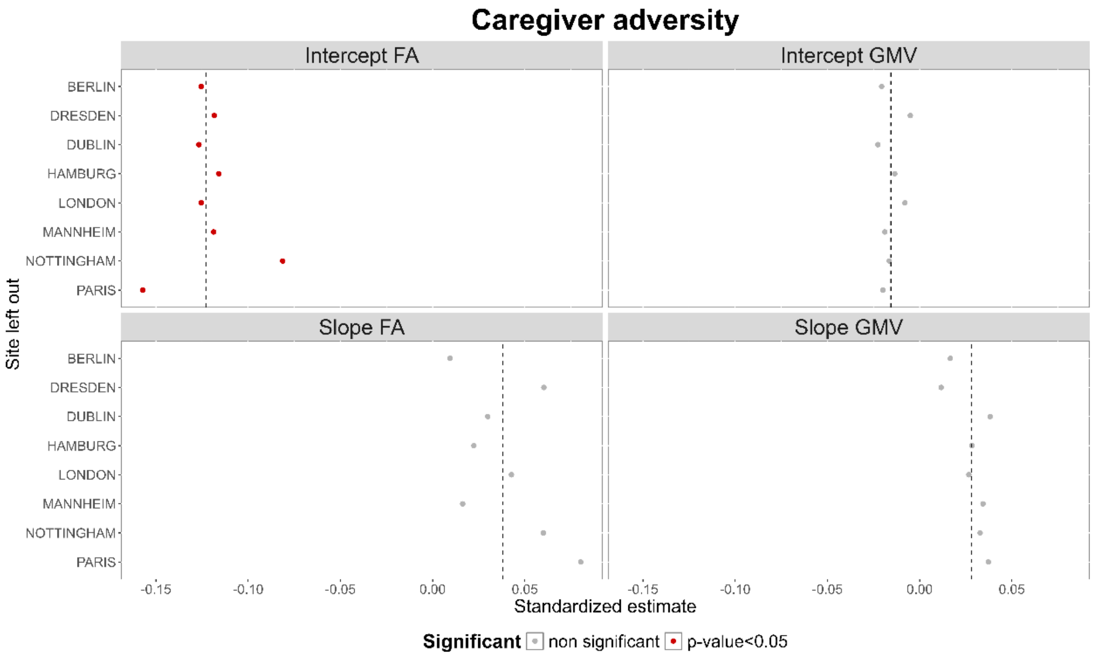

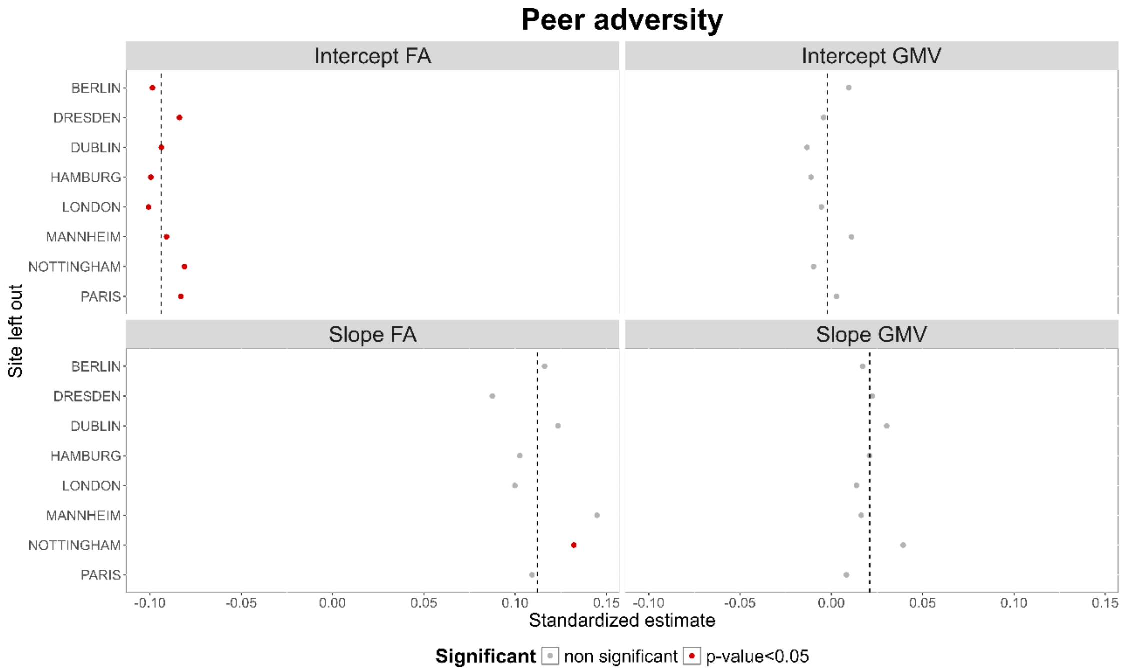

Peer adversity
Intercept FA
Intercept GMV
Slope FA
Slope GMV
Site left out
BERLIN
DRESDEN
DUBLIN
HAMBURG
LONDON
MANNHEIM
NOTTINGHAM
PARIS
-0.10
-0.05
0.00
0.05
0.10
0.15
Standardized estimate
Significant
non significant
p-value<0.05


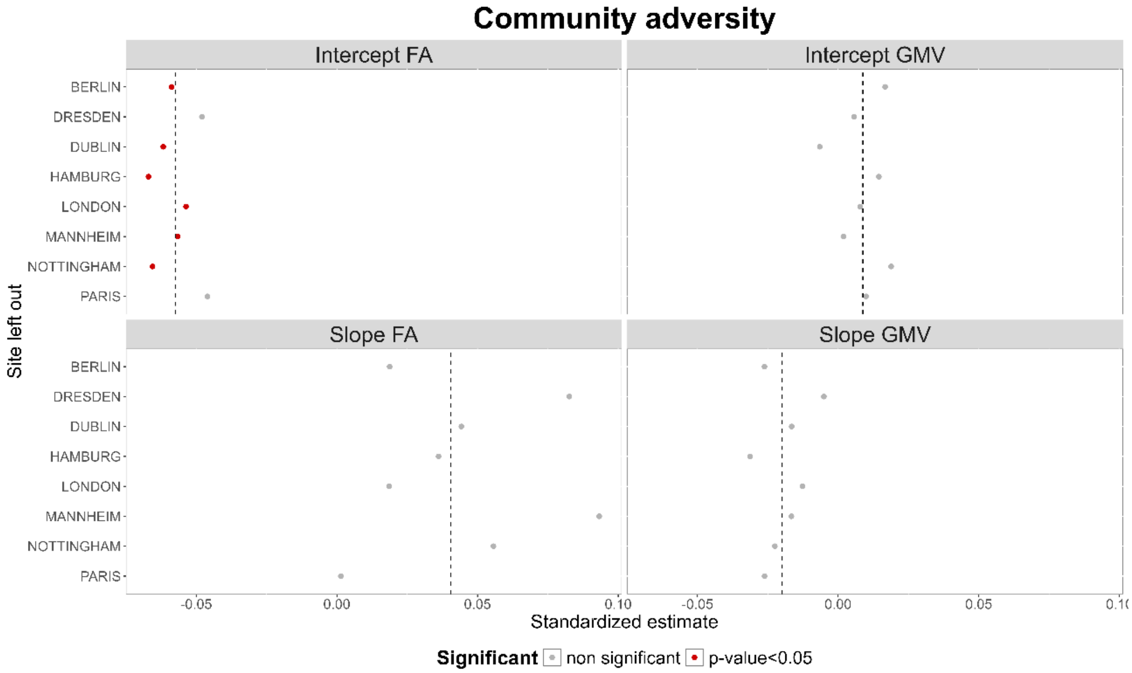

Community adversity
Intercept FA
Intercept GMV
Slope FA
Slope GMV
Site left out
BERLIN
DRESDEN
DUBLIN
HAMBURG
LONDON
MANNHEIM
NOTTINGHAM
PARIS
-0.05
0.00
0.05
0.10
Standardized estimate
Significant
non significant
p-value<0.05

### Distribution of adversity between males and females

#### ABCD

*Supplementary Figure 9. Distribution of the adversity components across males and females in ABCD*

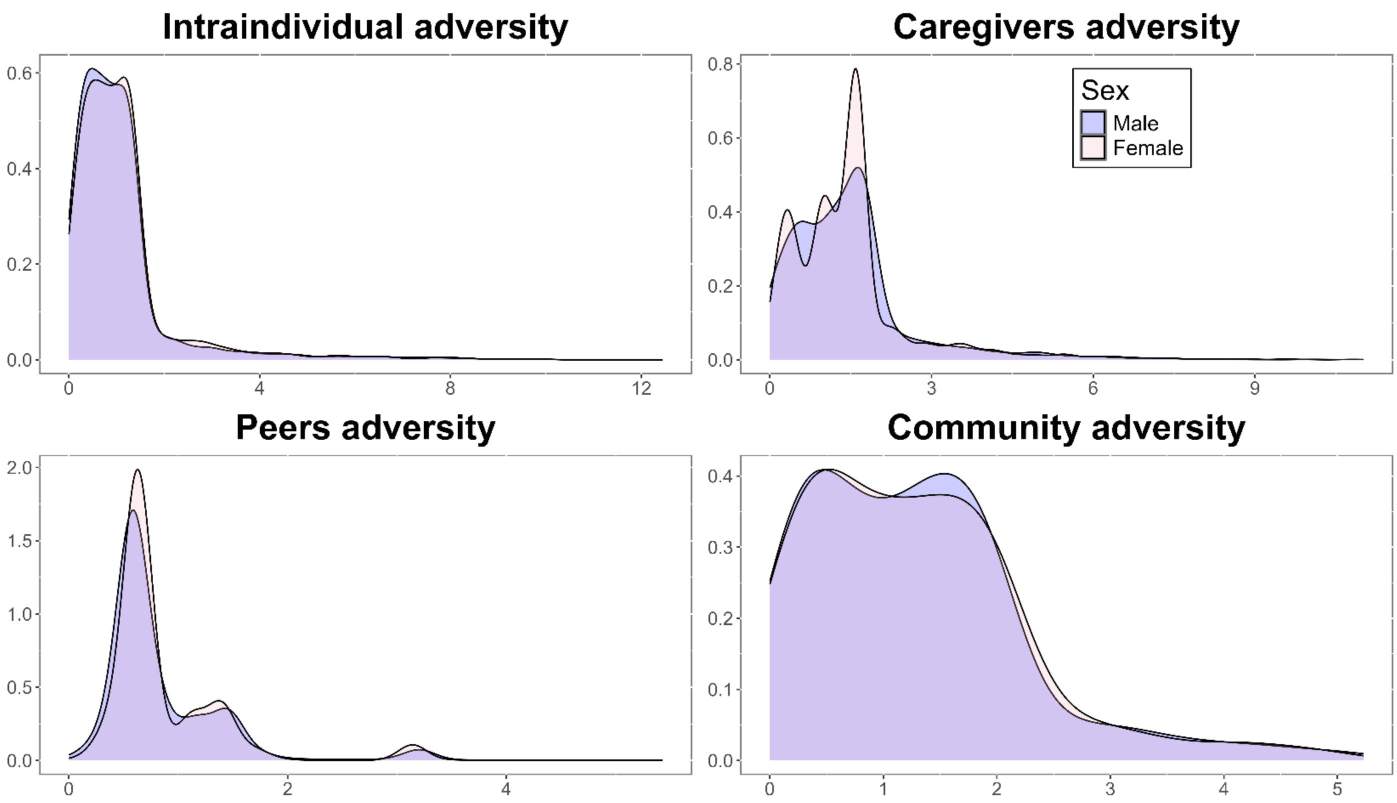


*Supplementary Figure 10. Distribution of the loadings for each of the adversity components across males and females in ABCD. The probabilistic component analyses were done separately for males and females.*

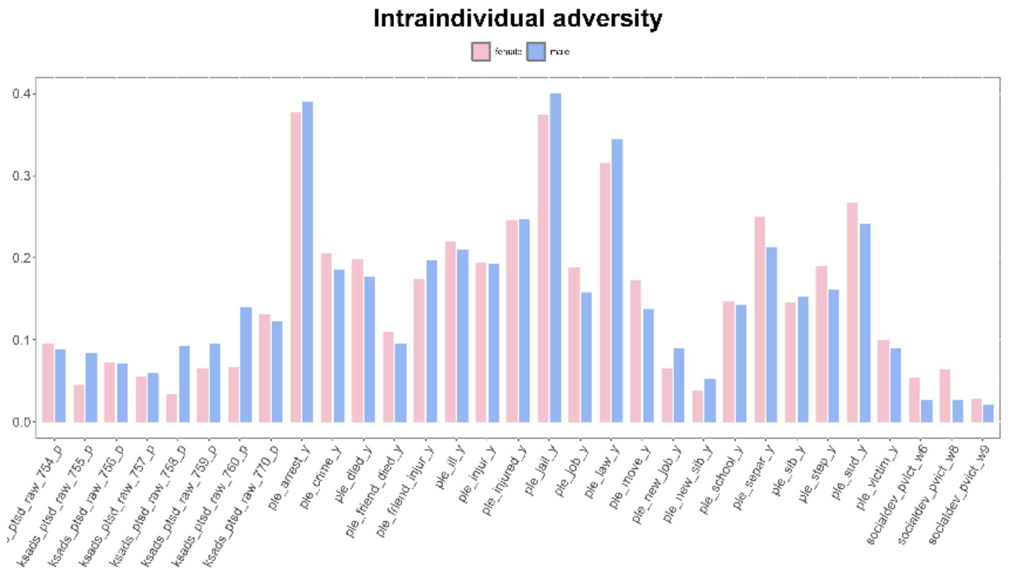
Intraindividual adversity
female
male
0.4
0.3
0.2
0.1
0.0
ksads_ptsd_raw_755_p
ksads_ptsd_raw_756_p
ksads_ptsd_raw_757_p
ksads_ptsd_raw_758_p
ksads_ptsd_raw_759_p
ksads_ptsd_raw_760_p
ksads_ptsd_raw_770_p
ple_arrest_y
ple_crime_y
ple_died_y
ple_friend_died_y
ple_friend_injur_y
ple_ill_y
ple_injur_y
ple_injured_y
ple_jail_y
ple_job_y
ple_law_y
ple_move_y
ple_new_job_y
ple_new_sib_y
ple_school_y
ple_separ_y
ple_sib_y
ple_step_y
ple_sud_y
ple_victim_y
socialdev_pvict_w6
socialdev_pvict_w8
socialdev_pvict_w9

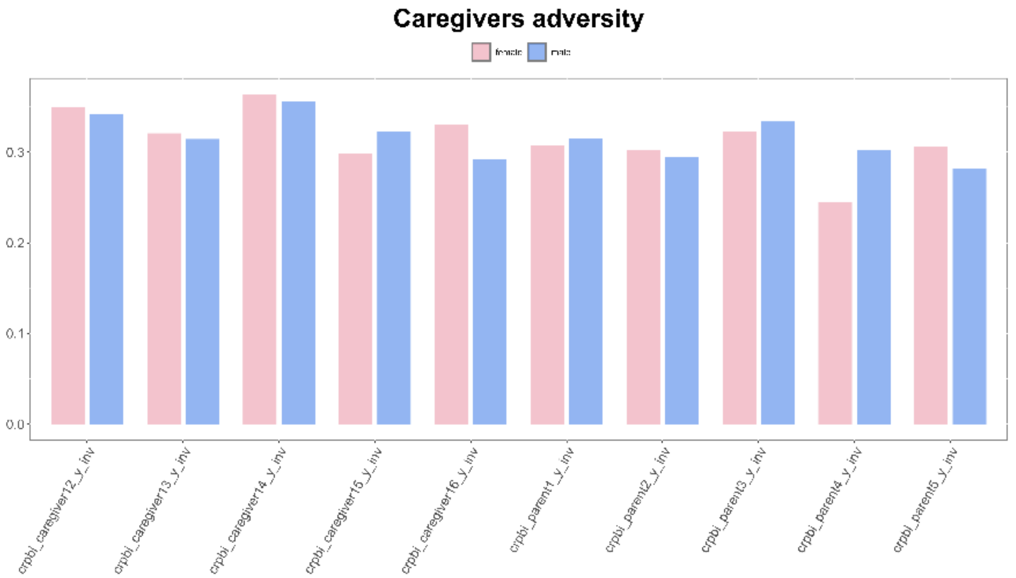
Caregivers adversity
female
male
0.3
0.2
0.1
0.0
crpbi_caregiver12_y_inv
crpbi_caregiver13_y_inv
crpbi_caregiver14_y_inv
crpbi_caregiver15_y_inv
crpbi_caregiver16_y_inv
crpbi_parent1_y_inv
crpbi_parent2_y_inv
crpbi_parent3_y_inv
crpbi_parent4_y_inv
crpbi_parent5_y_inv

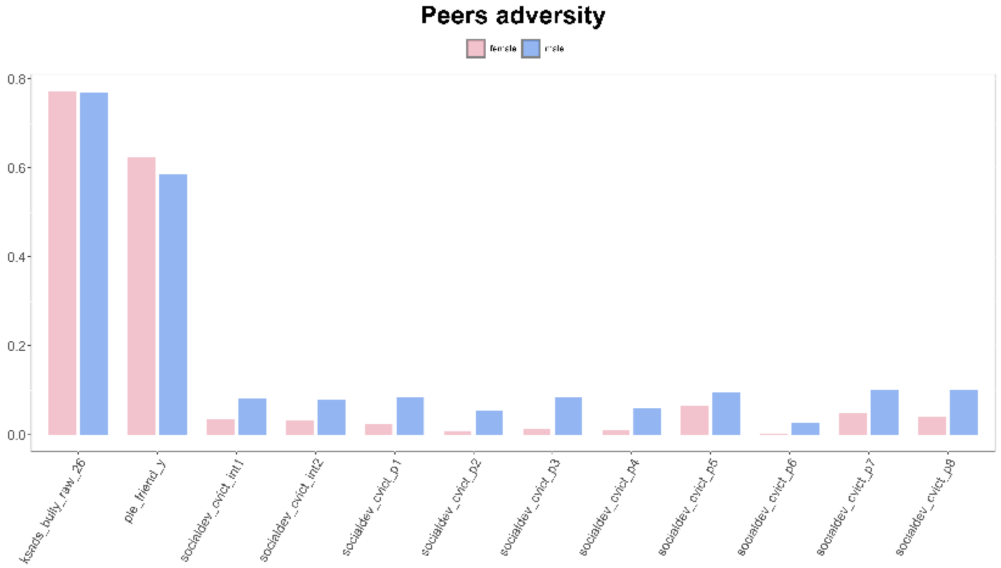
Peers adversity
female
male
0.8
0.6
0.4
0.2
0.0
ksads_bully_raw_26
ple_friend_y
socialdev_cvict_int1
socialdev_cvict_int2
socialdev_cvict_p1
socialdev_cvict_p2
socialdev_cvict_p3
socialdev_cvict_p4
socialdev_cvict_p5
socialdev_cvict_p6
socialdev_cvict_p7
socialdev_cvict_p8

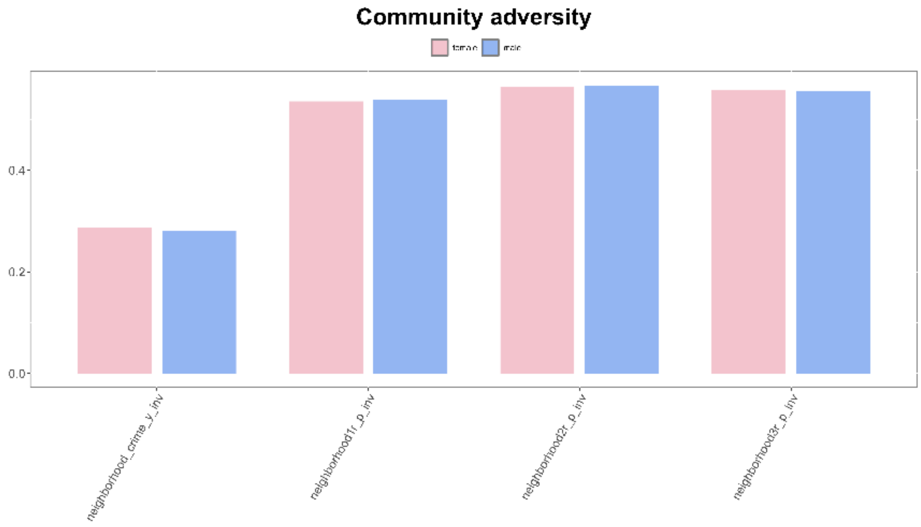


### IMAGEN

*Supplementary Figure 11. Distribution of the adversity components across males and females in IMAGEN*

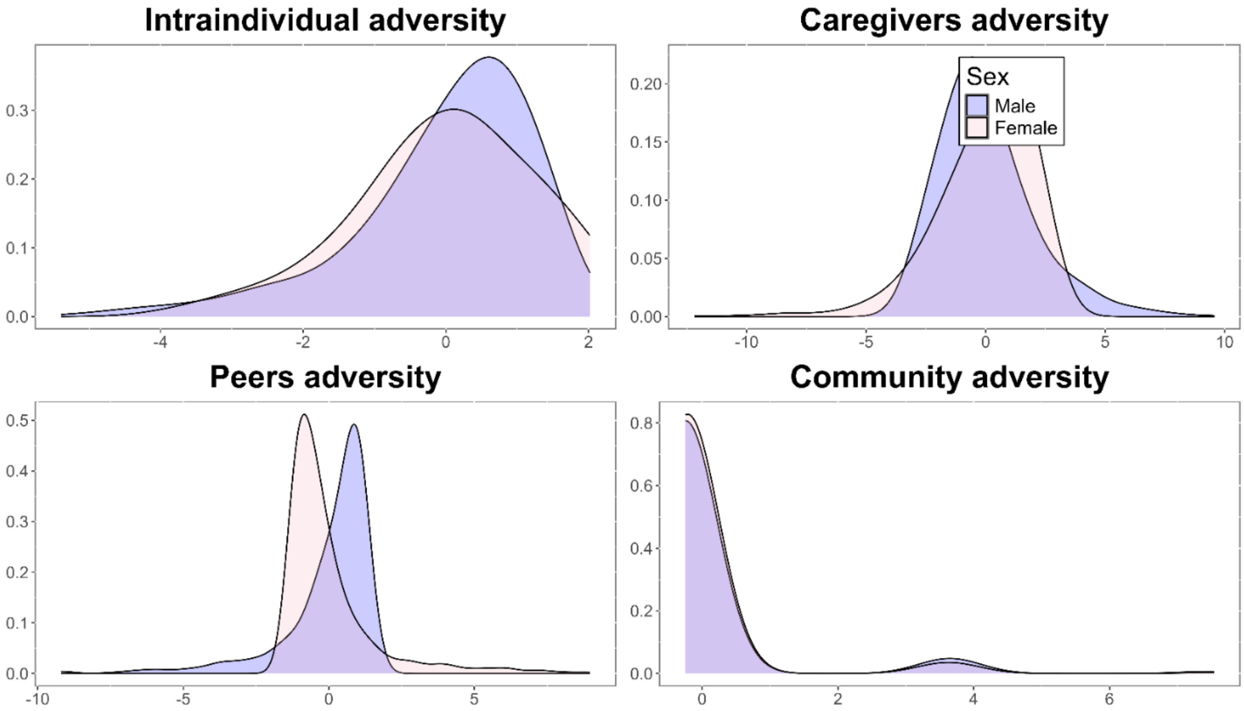

*Supplementary Figure 12. Distribution of the loadings for each of the adversity components across males and females in ABCD. The probabilistic component analyses were done separately for males and females.*

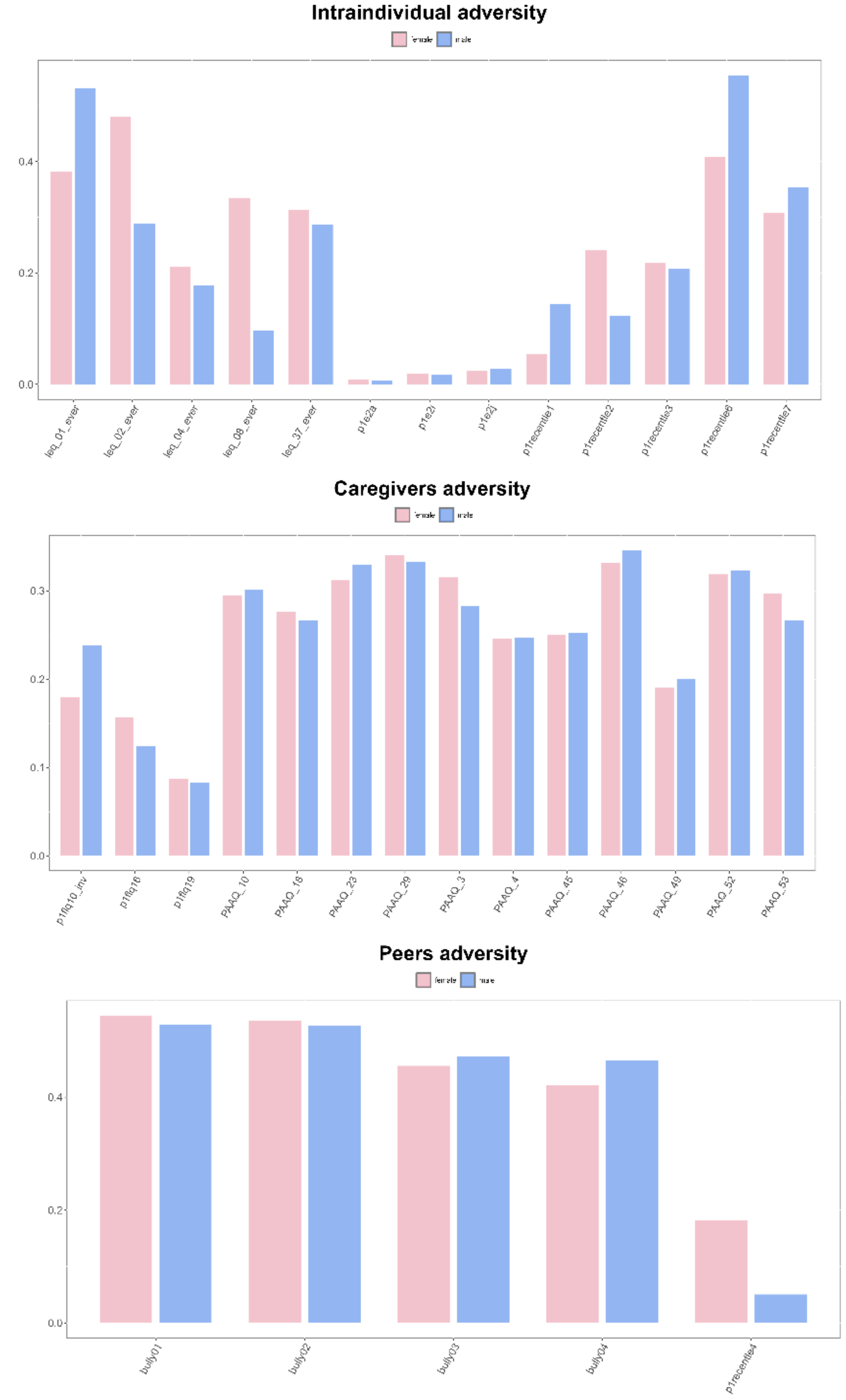

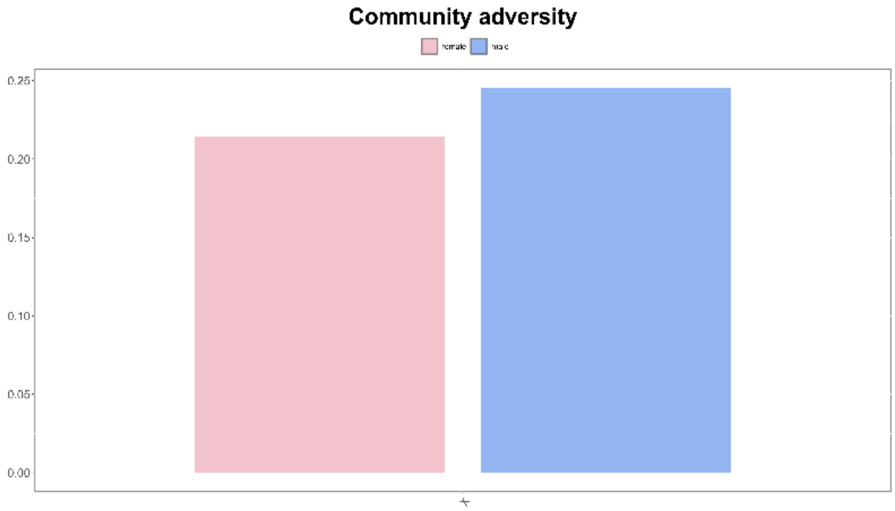
Community adversity
female
male
0.25
0.20
0.15
0.10
0.05
0.00

### Distribution of adversity between males and females

#### ABCD

*Supplementary Table 16. Parameters estimates of sex-stratified analysis for the bivariate latent growth curve model with the four levels of adversity predicting the intercepts and slopes of total cortical grey matter volume and mean fractional anisotropy in ABCD.*

**Full model with adversity in female (ABCD)**

*Fit indices: $x^2$ = 119.702, df= 34, p<0.001, RMSEA = 0.024, CFI= 0.996, TLI=0.990, SRMR= 0.009*

| Parameters | Estimate | Std.error | z-value | p-value | Std.estimate | CI lower | CI upper |
|---|---|---|---|---|---|---|---|
| **Intraindividual** | | | | | | | |
| Intercept GMV | -781.88 | 630.00 | -1.24 | 0.215 | -0.020 | -2017 | 453 |
| Intercept FA | -7.46E-05 | 2.71E-04 | -0.27 | 0.783 | -0.005 | -6.06E-04 | 4.57E-04 |
| Slope GMV | 164 | 352 | -2.51 | 0.641 | 0.021 | -525 | 853 |
| Slope FA | -2.05E-04 | 1.17E-04 | -1.75 | 0.079 | -0.071 | -4.35E-04 | 2.41E-05 |
| **Caregivers** | | | | | | | |
| Intercept GMV | -1805 | 676 | -2.67 | ***0.008*** | -0.041 | -3130 | -479 |
| Intercept FA | -7.12E-04 | 2.93E-04 | -2.43 | ***0.015*** | -0.040 | -1.29E-03 | -1.37E-04 |
| Slope GMV | -195 | 380 | -0.51 | 0.608 | -0.023 | -939 | 550 |
| Slope FA | 7.59E-05 | 1.42E-04 | 0.53 | 0.593 | 0.024 | -2.03E-04 | 3.54E-04 |
| **Peers** | | | | | | | |
| Intercept GMV | -3438 | 1367 | -2.51 | ***0.012*** | -0.040 | -6118 | -758 |
| Intercept FA | -2.98E-04 | 5.56E-04 | -0.54 | 0.591 | -0.009 | -1.39E-03 | 7.90E-04 |
| Slope GMV | 550 | 689 | 0.80 | 0.425 | 0.034 | -800 | 1899 |
| Slope FA | -1.13E-04 | 2.63E-04 | -0.43 | 0.667 | -0.018 | -6.28E-04 | 4.02E-04 |
| **Community** | | | | | | | |
| Intercept GMV | -5727 | 742 | -7.72 | ***<0.001*** | -0.116 | -7182 | -4273 |
| Intercept FA | -1.35E-03 | 3.09E-04 | -4.38 | ***<0.001*** | -0.068 | -1.96E-03 | -7.49E-04 |
| Slope GMV | -418 | 391 | -1.07 | 0.284 | -0.044 | -1184 | 347 |
| Slope FA | 4.52E-05 | 1.52E-04 | 0.30 | 0.766 | 0.013 | -2.53E-04 | 3.44E-04 |

**Full model with adversity in male (ABCD)**

| Parameters | Estimate | Std.error | z-value | p-value | Std.estimate | CI lower | CI upper |
|---|---|---|---|---|---|---|---|
| **Intraindividual** | | | | | | | |
| Intercept GMV | 14 | 649 | 0.02 | 0.982 | 0.000 | -1257 | 1286 |
| Intercept FA | -8.35E-05 | 2.79E-04 | -0.30 | 0.765 | -0.005 | -6.30E-04 | 4.63E-04 |
| Slope GMV | 80 | 271 | 0.30 | 0.767 | 0.011 | -451 | 611 |
| Slope FA | 1.46E-04 | 9.61E-05 | 1.52 | 0.130 | 0.061 | -4.27E-05 | 3.34E-04 |
| **Caregivers** | | | | | | | |
| Intercept GMV | -449 | 658 | -0.68 | 0.495 | -0.011 | -1738 | 841 |
| Intercept FA | -4.35E-04 | 2.38E-04 | -1.82 | 0.068 | -0.028 | -9.02E-04 | 3.27E-05 |
| Slope GMV | 724 | 254 | 2.85 | ***0.004*** | 0.113 | 226 | 1222 |
| Slope FA | 1.89E-05 | 1.14E-04 | 0.17 | 0.869 | 0.009 | -2.05E-04 | 2.43E-04 |
| **Peers** | | | | | | | |
| Intercept GMV | -3712 | 1373 | -2.70 | ***0.004*** | -0.039 | -6403 | -1020 |
| Intercept FA | -7.42E-05 | 5.43E-04 | -0.14 | 0.891 | -0.002 | -1.14E-03 | 9.90E-04 |
| Slope GMV | -138 | 654 | -0.21 | 0.832 | -0.010 | -1421 | 1144 |
| Slope FA | 2.53E-04 | 2.71E-04 | 0.93 | 0.350 | 0.052 | -2.78E-04 | 7.83E-04 |
| **Community** | | | | | | | |
| Intercept GMV | -6182 | 781 | -7.91 | ***<0.001*** | -0.115 | -7713 | -4651 |
| Intercept FA | -1.23E-03 | 3.01E-04 | -4.10 | ***<0.001*** | -0.061 | -1.83E-03 | -6.44E-04 |
| Slope GMV | 350 | 371 | 0.95 | 0.345 | 0.042 | -376 | 1076 |
| Slope FA | 3.31E-04 | 1.49E-04 | 2.22 | ***0.004*** | 0.119 | 3.84E-05 | 6.24E-04 |

### IMAGEN

*Supplementary Table 17. Parameters estimates of sex-stratified analysis for the bivariate latent growth curve model with the four levels of adversity predicting the intercepts and slopes of total cortical grey matter volume and mean fractional anisotropy in IMAGEN.*

**Full model with adversity in female (IMAGEN)**

*Fit indices: $x^2$ = 91.441, df= 34, p<0.001, RMSEA = 0.060, CFI= 0.988, TLI=0.973, SRMR= 0.019*

| Parameters | Estimate | Std.error | z-value | p-value | Std.estimate | CI lower | CI upper |
|---|---|---|---|---|---|---|---|
| **Intraindividual** | | | | | | | |
| Intercept GMV | -1645 | 210 | -0.78 | 0.434 | -0.039 | -5769.36 | 2478.85 |
| Intercept FA | -6.00E-05 | 5.25E-04 | -0.11 | 0.909 | -0.004 | -1.09E-03 | 9.70E-04 |
| Slope GMV | 2933 | 1687 | 1.74 | 0.082 | 0.091 | -374 | 6240 |
| Slope FA | 7.42E-04 | 4.39E-04 | 1.69 | 0.091 | 0.140 | -1.19E-04 | 1.60E-03 |
| **Caregivers** | | | | | | | |
| Intercept GMV | -1287 | 1223 | -1.05 | 0.293 | -0.048 | -3685 | 1110 |
| Intercept FA | -1.61E-03 | 3.66E-04 | -4.41 | ***<0.001*** | -0.174 | -2.33E-03 | -8.97E-04 |
| Slope GMV | 486 | 851 | 0.57 | 0.568 | 0.024 | -1183 | 2155 |
| Slope FA | 9.23E-05 | 2.16E-04 | 0.43 | 0.668 | 0.027 | -3.30E-04 | 5.15E-04 |
| **Peers** | | | | | | | |
| Intercept GMV | 1029 | 1507 | 0.68 | 0.494 | 0.031 | -1924 | 3982 |
| Intercept FA | -7.99E-04 | 4.04E-04 | -1.98 | ***0.048*** | -0.071 | -1.59E-03 | -7.53E-06 |
| Slope GMV | 379 | 1032 | 0.37 | 0.714 | 0.015 | -1644 | 2402 |
| Slope FA | 2.16E-04 | 3.58E-04 | 0.60 | 0.545 | 0.053 | -4.84E-04 | 9.17E-04 |
| **Community** | | | | | | | |
| Intercept GMV | 1070 | 2118 | 0.51 | 0.613 | 0.020 | -3082 | 5222 |
| Intercept FA | -4.47E-04 | 5.09E-04 | -0.88 | 0.380 | -0.025 | -1.44E-03 | 5.50E-04 |
| Slope GMV | -175 | 1505 | -0.12 | 0.908 | -0.004 | -3125 | 2776 |
| Slope FA | 1.66E-04 | 4.18E-04 | 0.40 | 0.690 | 0.025 | -6.52E-04 | 9.85E-04 |

**Full model with adversity in male (IMAGEN)**

| Parameters | Estimate | Std.error | z-value | p-value | Std.estimate | CI lower | CI upper |
|---|---|---|---|---|---|---|---|
| **Intraindividual** | | | | | | | |

| | | | | | | | |
|---|---|---|---|---|---|---|---|
| Intercept GMV | -3854 | 2975 | -1.30 | 0.195 | -0.075 | -9684 | 1976 |
| Intercept FA | 1.15E-03 | 5.56E-04 | 2.06 | ***0.039*** | 0.076 | 5.82E-05 | 2.24E-03 |
| Slope GMV | 2162 | 2779 | 0.78 | 0.437 | 0.054 | -3285 | 7609 |
| Slope FA | -4.38E-04 | 4.03E-04 | -1.09 | 0.276 | -0.140 | -1.23E-03 | 3.51E-04 |
| **Caregivers** | | | | | | | |
| Intercept GMV | -3077 | 1189 | -2.59 | ***0.010*** | -0.097 | -5408 | -746 |
| Intercept FA | -8.27E-04 | 4.03E-04 | -2.05 | ***0.040*** | -0.089 | -1.62E-03 | -3.72E-05 |
| Slope GMV | 638 | 952 | 0.67 | 0.503 | 0.025 | -1229 | 2504 |
| Slope FA | -1.42E-04 | 2.41E-04 | -0.59 | 0.555 | -0.073 | -6.13E-04 | 3.30E-04 |
| **Peers** | | | | | | | |
| Intercept GMV | 1601 | 2274 | 0.70 | 0.481 | 0.039 | -2856 | 6058 |
| Intercept FA | -1.45E-03 | 4.37E-04 | -3.33 | ***0.001*** | -0.120 | -2.31E-03 | -5.97E-04 |
| Slope GMV | 575 | 1855 | 0.31 | 0.757 | 0.018 | -3060 | 4210 |
| Slope FA | -1.45E-03 | 3.75E-04 | 2.17 | ***0.030*** | 0.323 | 7.78E-05 | 1.55E-03 |
| **Community** | | | | | | | |
| Intercept GMV | -2499 | 1815 | -1.38 | 0.169 | -0.039 | -6057 | 1058 |
| Intercept FA | -1.66E-03 | 7.44E-04 | -2.23 | ***0.026*** | -0.088 | -3.12E-03 | -2.02E-04 |
| Slope GMV | -1519 | 1531 | -0.99 | 0.321 | -0.030 | -4520 | 1482 |
| Slope FA | 2.29E-04 | 4.30E-04 | 0.53 | 0.595 | 0.058 | -6.14E-04 | 1.07E-03 |
